# 3D Biomimetic Liver Cancer Model: Diethylnitrosamine-Induced Proteomic Dysregulations in Stromal-Epithelial Milieu

**DOI:** 10.1101/2025.05.17.654447

**Authors:** Salmma S Salihah, Bareerah Bibi, Sehrish Khan, Muhammad Tahir, Sana Mahmood, Massoud Vosough, Agnese Razzoli, Simone Sidoli, Asma Gul

## Abstract

Hydrogel-based three-dimensional (3D) co-culture systems are emerging as biomimetic platforms that more accurately recapitulate tissue architecture and microenvironmental interactions compared to conventional two-dimensional (2D) cultures. This study introduces an engineered 3D liver-like model to investigate compartment-specific responses to the potent hepatocarcinogen Diethylnitrosamine (DEN), with a focus on early events in carcinogenesis and tumor-stroma interactions. AML12 and 3T3 cell lines were treated with DEN or vehicle either in 2D culture or in 3D hydrogels in four experimental groups: (1) DEN-treated AML12 with vehicle-treated 3T3, (2) DEN-treated 3T3 with vehicle-treated AML12, (3) both cell types DEN-treated, and (4) both vehicle-treated. The cultured recombinants were subjected to proteomic profiling via mass spectrometry, followed by bioinformatics analysis and the results were validated through immunocytochemical staining (ICC). Gene ontology analysis revealed that cytoskeletal, RNA metabolism, and scaffold/adaptor proteins were among the most significantly enriched in 3D versus 2D models. Structural proteins emerged exclusively in mixed 3D co-cultures, reinforcing the organotypic nature of the system. Enriched pathways in 3D included intermediate filament organization, actin dynamics, and focal adhesion—pathways closely associated with liver carcinogenesis. Protein-protein interaction analysis demonstrated maximal network complexity in 3D cultures where both compartments were DEN-exposed. Survival analysis further identified poor-prognosis biomarkers (KRT20, KRT15, KRT14) uniquely enriched in this condition. ICC staining supported the proteomic findings. This organoid-like 3D co-culture model provides a physiologically relevant platform for investigating early-stage liver carcinogenesis and highlights the critical role of stromal–epithelial interactions. Its ability to replicate organ-level complexity and generate clinically relevant proteomic signatures supports its utility in translational cancer research and future drug discovery applications.

## 1. Introduction

The traditional 2D culture systems have long been used in cancer research but cannot ideally mimic physiological conditions including inter- and intra-tumor heterogeneity and the mechanisms involved in the initiation of cancer due to the lack of cell-cell communication and cell-matrix interactions [1,2]. Alternatively, 3D cultures support the acquisition and maintenance of a phenotype closer to the naturalistic phenotype and gene expression closer to the in vivo tumors [3]. Moreover, 3D co-culture of hepatocyte and non-parenchyma cells has the potential to represent in vivo liver activity in diverse functional states [4].

Conventionally hepatocellular carcinoma (HCC) is assumed to originate from carcinogen acting on the epithelial cells [5] while stromal cells inhibit HCC by restraining cellular proliferation in normal tissue and in the initial stages of carcinogenesis [6,7]. In the later stages of HCC, malignant epithelial cells reprogram the tumor microenvironment (TME), and stromal cells become not just active bystanders in the pathogenesis of HCC [8], but they are transformed into cancer-associated fibroblasts (CAF) through paracrine signaling by the tumor and start playing an important role in tumor initiation and progression [9]. An alternative approach to the initiation of carcinogenesis revealed that stroma is the target of nitrosomethyl urea (NMU) in memory gland carcinogenesis and hence postulated that altered tissue architecture is at the core of carcinogenesis. Objectively, however the specific target(s) upon which carcinogenic agents act to carry out neoplastic transformation in various tissues are yet to be identified [10].

Several studies employing three-dimensional (3D) liver models have revealed stromal changes induced by toxicants or fibrogenic stimuli, even when diethylnitrosamine (DEN) was not directly tested. For instance, 3D hepatic co-cultures and spheroid models have demonstrated that genotoxic agents can provoke extracellular matrix (ECM) remodeling, activate fibroblastic components, and disrupt epithelial-stromal crosstalk, phenomena that remain largely undetectable in traditional two-dimensional (2D) cultures [11,12]. These studies underscore the unique capacity of 3D systems to capture cell-matrix and paracrine signaling events critical to early carcinogenic processes. Like chemotherapeutics and fibrogenic agents, DEN—a classical DNA-alkylating hepatocarcinogen—likely alters ECM-cell interactions, thereby contributing to the initiation of a tumor-permissive microenvironment. Such disruptions are detectable only in physiologically relevant 3D contexts, where spatial and mechanical cues are preserved [13]. Building on this conceptual framework, our spatial proteomics approach introduces a novel layer by revealing zonated stromal-epithelial dysregulation following DEN exposure. Unlike prior 3D studies that described global micro environmental shifts, our work dissects hepatic zone-specific protein alterations, offering new mechanistic insight into how regional stromal remodeling contributes to DEN-driven hepatocarcinogenesis.

To reflect the target of hepatic carcinogens, a physiologically relevant model needs to be developed and characterized. Only 20-50% of the transcribed genes are expressed into functional proteins [3] and proteins are the final effectors of cellular machinery. Owing to the fact that the proteomic changes often get overlooked as the cells arrange themselves into complex 3D structures, we carried out proteomic characterization of 3D co-culture model in comparison with the traditional 2D model. The superior 3D model was then used to compare proteomic variation in response to Diethylnitrosamine (DEN) (a potent liver carcinogen) in four different experimental groups designed to identify the target of carcinogen and the overall protein expression changes arising in response to the hepatic carcinogen. The results were validated through immunocytochemistry. The findings of this study are expected to provide insightful information about the identification of effective therapeutic targets and drug discovery regimes.

## 2. Materials and Methods

### 2.1. Culture Maintenance

AML12 (Alpha mouse liver 12, CRL-2254) and 3T3 (mouse embryonic fibroblasts, CRL-1658) cell lines were obtained from ATCC and cultured in 25 cm^3^ flasks (Corning, USA) according to the manufacturer’s instructions. The base medium used for AML12 was DMEM: F12 (Gibco) whereas 3T3 utilized DMEM (Gibco) only. Both mediums were supplemented with 10% FBS, 5 ng/ml selenium, 10 µg/ml insulin, 5.5 µg/ml Transferrin, and 40 ng/ml dexamethasone. For co-culture experiments, mediums for both cell types were mixed in 1:1 and tested on each cell line alone before being used in co-culture experiments to ensure appropriate growth and behavior of cells. The culture medium was replaced every two days for all cell types. All cultures were incubated at 37 LJC and 5% carbon dioxide.

### 2.2. Carcinogen Exposure and Recombinant Culture

2D cultures of AML12 and 3T3 cells were separately exposed to Diethylnitrosamine (CAS # 55-18-5, Sigma) at a concentration of 0.2 mg/ml for 4 hours. DEN was withdrawn after 4 hours and cells were rinsed thrice with 1X PBS. Following three consecutive exposures, each separated by an interval of 1 week, the cells were recombined into four distinct groups, keeping the ratio of 3T3 to AML12 0.15:1 which mimics the approximate physiological ratio of stromal to epithelial cells in the liver. Group 1 (cAML) consisted of epithelial cells (AML12) exposed to DEN and stromal cells (3T3) exposed to vehicle (normal saline). Group 2 (c3T3) consisted of AML12 exposed to vehicle and 3T3 exposed to DEN. In Group 3 (cBoth) AML12 and 3T3 both were exposed to DEN whereas in group 4 (An3n) both cell types were exposed to vehicle. Recombinants were then grown as adherent 2D (as described above) and 3D cultures. For 3D culture, cells were suspended in 1mL rat tail collagen and seeded in 24 well plates. The gels were allowed to solidify for 30 min at 37 LJC and 5% carbon dioxide. Culture medium (1mL) was added to the solidified gels and the gels were detached from the walls and the bottom of 24 well plates using a pipette tip. Cultures were maintained for 2 weeks, and the medium was changed every 2-3 days.

### 2.3. Sample Preparation for Mass Spectrometry

Cells in the 2D culture were detached with 2ml trypsin and centrifuged to obtain the cell pellet. A protease phosphatase inhibitor cocktail from Sigma (cat # PPC1010) was added freshly to RIPA buffer (1ml per 100 ml of extraction buffer) and added to the cell pellet.

The 3D gels were harvested and rinsed briefly with phosphate buffer saline (PBS). Afterwards, 2 mL of 0.125% collagenase (Fischer Scientific) was added to digest the collagen gel and release the cells in the culture medium. The harvested cells were washed with PBS and centrifuged to get the pellet. The pellet was homogenized in 250µl of RIPA buffer and freshly added mix of phosphatase inhibitor 100X (Sigma Aldrich). The lysate was centrifuged at 14000 rpm and 4 LJC. The supernatant was collected and quantified. Protein estimation for the samples from both 2D and 3D culture was carried out through Bradford Assayas described by Bradford [10].

### 2.4. S-trap protein digestion

Disulfide bond reduction was carried out by resuspending the cell pellet in 100 µl buffer (50 mM ammonium bicarbonate (pH = 8), 5 mM DTT, 5% SDS) for 1 hour followed by alkylation with 20 mM iodoacetamide for 30 minutes, in the dark. A final concentration of 1.2% phosphoric acid was then added to the sample. Samples were diluted in six volumes of binding buffer (10 mM ammonium bicarbonate and 90% methanol). The protein solution was gently mixed and loaded to a S-trap filter (Protifi) and centrifuged for 30 sec at 500g. After washing the samples twice with binding buffer, 1 µg of sequencing grade trypsin (Promega) was diluted in 50 mM ammonium bicarbonate and added into the S-trap filter and samples for digestion for 18 h at 37°C. Peptides were eluted in three steps: (i) 40 µl of 50 mM ammonium bicarbonate, (ii) 40 µl of 0.1% TFA and (iii) 40 µl of 60% acetonitrile and 0.1% TFA. The peptide solution was pooled and centrifuged for 30 sec at 1,000 g and then dried in a vacuum centrifuge.

### 2.5. Sample desalting

Prior to mass spectrometry analysis, samples were desalted using a 96-well plate filter (Orochem) packed with 1 mg of Oasis HLB C-18 resin (Waters). Briefly, the samples were resuspended in 100 µl of 0.1% TFA and loaded onto the HLB resin, which was previously equilibrated using 100 µl of the same buffer. After washing with 100 µl of 0.1% TFA, the samples were eluted with a buffer containing 70 µl of 60% acetonitrile and 0.1% TFA and then dried in a vacuum centrifuge.

### 2.6. LC-MS/MS Acquisition and Analysis

Protein samples were resuspended in 10 µl of 0.1% TFA and loaded onto a Dionex RSLC Ultimate 300 (Thermo Scientific), coupled online with an Orbitrap Fusion Lumos (Thermo Scientific). A two-column system was used for chromatographic separation, consisting of a C-18 trap cartridge (300 µm ID, 5 mm length) and a picofrit analytical column (75 µm ID, 25 cm length) packed in-house with reversed-phase Repro-Sil Pur C18-AQ 3 µm resin. Peptides were separated using a 60 min gradient from 4-30% buffer B (buffer A: 0.1% formic acid, buffer B: 80% acetonitrile + 0.1% formic acid) at a flow rate of 300 nl/min. The mass spectrometer was set to acquire spectra in a data-dependent acquisition (DDA) mode. Briefly, the full MS scan was set to 300-1200 m/z in the orbitrap with a resolution of 120,000 (at 200 m/z) and an AGC target of 5×10e5. MS/MS was performed in the ion trap using the top speed mode (2 secs), an AGC target of 1×10e4 and an HCD collision energy of 35.

Proteome Discoverer software (v2.5, Thermo Scientific) using SEQUEST search engine and the SwissProt mouse database (updated March 2024) were used to search proteome raw file. The search for total proteome included fixed modification of carbamidomethyl cysteine and variable modification of N-terminal acetylation. Trypsin with the allowance of two missed cleavages was specified as the digestive enzyme. Mass tolerance was set to 0.2 Da for product ions 10 pm for and precursor ions.

### 2.7. Statistical Data Analysis

The statistical analysis of the data was performed by importing protein groups from MaxQuant. The analysis was carried out on Perseus v.1.6.10.50 after removing potential contaminants, reverse database hits and protein identified by site, log2 transformation was carried out. The samples were initially categorized into two groups depending on the culture type and then further into four groups depending on the treatments. Proteins were further filtered to include only those with at least three valid values in each group. Significance testing of protein LFQ intensities was carried out by post hoc Tukey’s HSD test for oneway ANOVA with a p-value threshold of <0.05 (Benjamini-Hochberg FDR at 0.05) considered significant. Groupwise comparisons were carried out using functional analysis tool. Principal component analysis was carried out to see the differentiation of treated and control samples. Heatmaps were generated using Perseus v.1.6.10.50, displaying protein intensities with non-supervised hierarchical clustering of rows. Volcano plots were made to represent differentially expressed proteins.

### 2.8. Bioinformatics Analysis

Overrepresentation analysis was performed using the *clusterProfiler* package in R to assess the statistical significance of the enrichment. Additionally, the PANTHER classification system was employed to identify and categorize the protein classes enriched in each group, providing further insights into the functional roles of the DEPs within the biological context. Functional associations or physical interactions between protein networks was visualized using STRING database and Cytoscape. The significant module for each network was obtained by MCODE using default parameters. The clinical value of differentially expressed proteins was estimated by converting 20 hub proteins to their human homologs using the “biomaRt” R package. Protein expression of human homologous proteins corresponding to DEPs was matched with overall survival using the UALCAN tool.

### 2.9. Immunocytochemistry

The gels sitting in 10% phosphate-buffered formalin were washed with PBS and changed to 70% ethanol and stained with Carmine Alum overnight following a protocol described previously [14]. After staining, the whole mounts were dehydrated in 70%, 95% and 100% ethanol, cleared in xylene and mounted with PermountTM (Fisher Scientific, Atlanta, GA).

An antigen-retrieval method based on microwave pretreatment and 0.01 M sodium citrate buffer (pH 6) was used as previously described [14]. Antibodies, including anti Ki67 antibody (Cat # PA0118, Leica biosystems), Vimentin (Cat # NCL-VIM-V9, Leica biosystems) and E-Cadherin (Cat# PA0387, Leica biosystems) were used at 1:400, 1:1000, 1:1000-4000 dilutions, respectively. The antigen-antibody reaction was visualized using the streptavidin-peroxidase complex, with diaminobenzidine tetrahydrochloride (Sigma-Aldrich) as the chromogen. Counterstaining was carried out with Harris’ hematoxylin. Images were captured with an Olympus digital camera attached to an Olympus BX53 microscope. In this section, where applicable, authors are required to disclose details of how generative artificial intelligence (GenAI) has been used in this paper (e.g., to generate text, data, or graphics, or to assist in study design, data collection, analysis, or interpretation). The use of GenAI for superficial text editing (e.g., grammar, spelling, punctuation, and formatting) does not need to be declared.

## 3. Results

### 3.1. Identification of Deferentially Expressed Proteins (DEP)

In this study, protein samples isolated from cell cultures were analyzed through label free LC-MS which led to the confident identification and relative quantification of total 5367 proteins shared across all the biological replicates within experimental groups. After the filtration of potential contaminants, only identified by site and reverse data base hits, 3972 proteins were selected and divided into 4 groups (i) 3T3 treated with DEN and AML 12 treated with vehicle (c3T3), (ii) AML 12 treated with DEN and 3T3 treated with vehicle (cAML), (iii) Both AML12 and 3T3 treated with DEN (cBoth), (iv) both AML 12 and 3T3 treated with vehicle (An3n/Ctrl). From this point onwards separate Log2 transformation of LFQ intensities of proteins in 2D and 3D groups was carried out. Both groups of 2D and 3D were separately subjected to filtration based on 70% values in each group reducing the total number of proteins to 3921 in 2D and 3827 in 3D. After filtration, post Hoc Tukey’s HSD test for one way ANOVA was used to test the significance of the expressed protein based on the difference between LFQ intensities of treatment and control groups. In 2D culture, 20 proteins were found to be common between treatment and control groups whereas 12, 64 and 85 proteins were significantly differentially expressed in c3T3, cBoth and cAML respectively. Among the differentially expressed proteins in cAML, 47 proteins were up regulated, and 40 proteins were downregulated, in c3T3 26 protein were upregulated and 40 proteins were downregulated, in cBoth group 29 proteins were upregulated and 37 proteins were downregulated. In 3D culture, 5 proteins were found to be common between treatment and control groups whereas 42, 79 and 38 proteins were significantly differentially expressed in c3T3, cBoth and cAML12 respectively. Among the differentially expressed proteins in cAML 17 proteins were upregulated and 23 proteins were downregulated, in c3T3 31 proteins were upregulated and 13 proteins were downregulated, 39 proteins were upregulated whereas 42 proteins were downregulated in cBoth group. Supplementary Table 1 represents the protein IDs, gene names and unique peptides of the perturbed proteins in treatment and control groups.

Venn Diagrams of 2D indicate that 50, 25 and 16 proteins were uniquely differentially expressed in cAML, c3T3 and cBoth respectively. In 3D culture, 29, 24 and 59 proteins were uniquely differentially expressed in cAML, c3T3 and cBoth (Figure 1 a and b).

**Figure 1:**
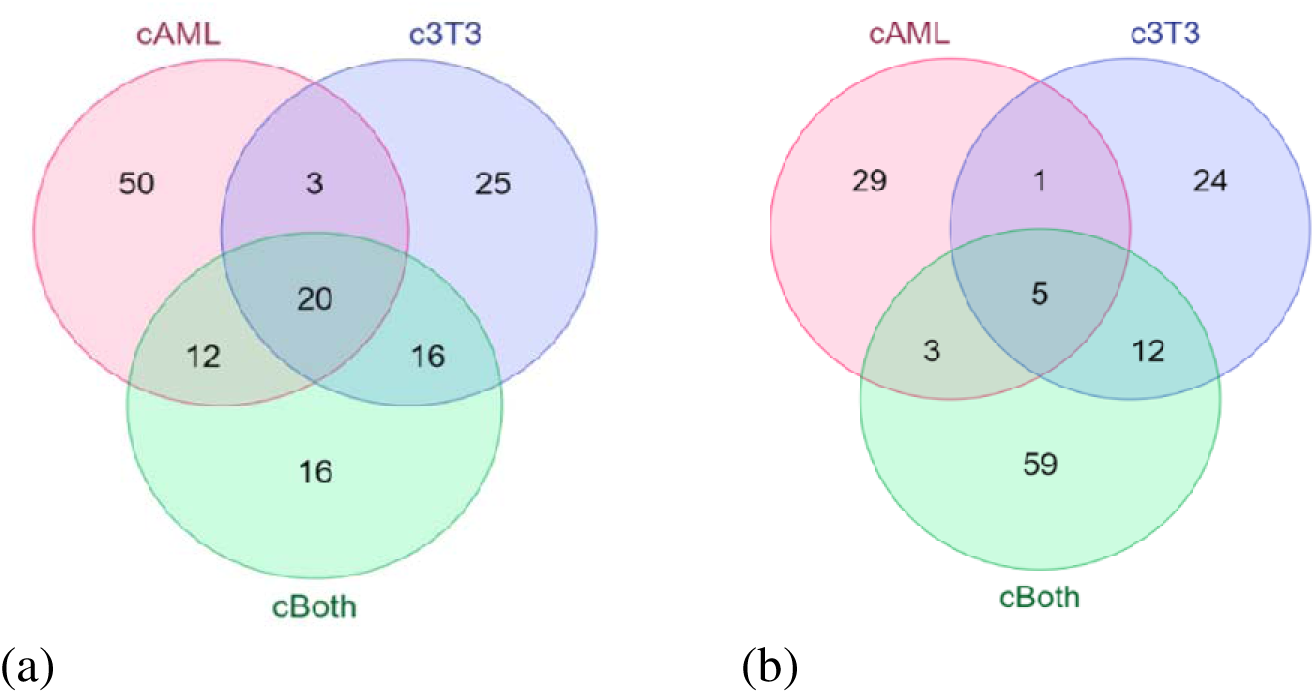
Differentially expressed and overlapping proteins (**a**) Ven diagram representing differential & common proteins in 2D; (**b**) Ven diagram representing differential and common proteins in 3D.

Heatmap of 2D represents heierarchical clustring of different treatment groups and comparsion between different replicates in each group (Figure 2a). Principal component analysis of 2D shows distinct spatial clustring with 36.7% contribution of PC1 and 16.5% contribution of PC2 to the variance (Figure 2B). Heatmap of 3D group is represented in figure 2C. PCA analysis of 3D shows that PC1 contributes 29.9% to the variance whereas PC2 contributes 20.1% to th variance (Figure 2D). Volcano plots were generated to determine the number of differentially expressed proteins for each group using a two-sided t-test with an FDR of 0.05 and mean - log2(x) fold change of ±1.0 (Figure 3 a-f).

**Figure 2:**
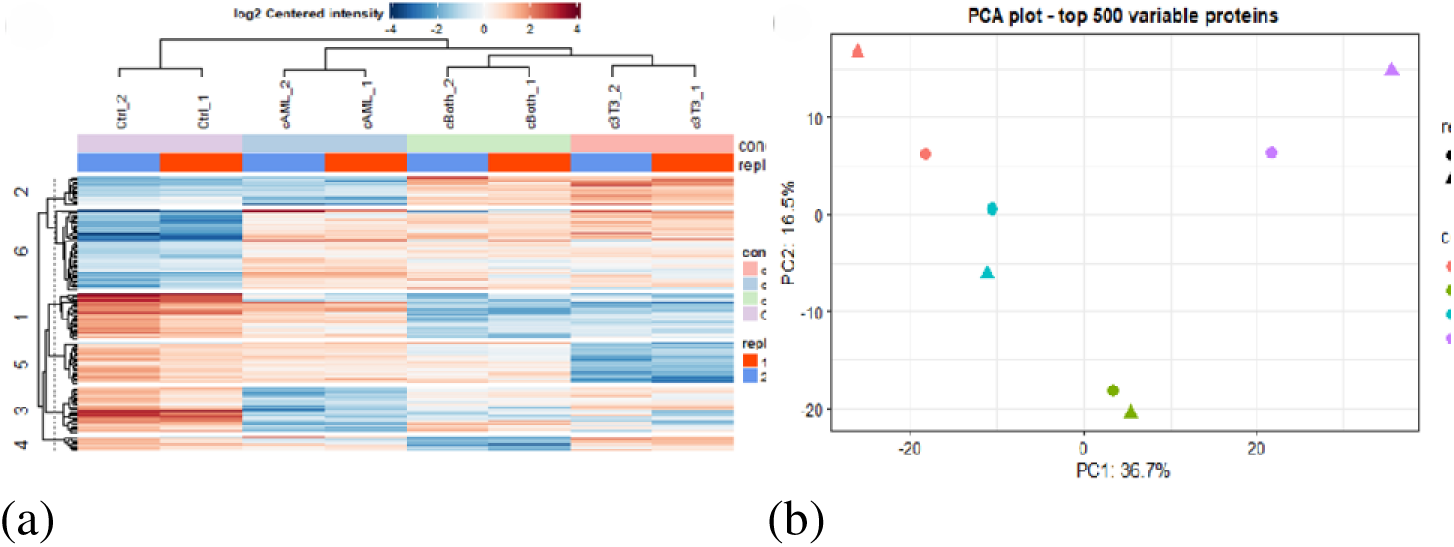
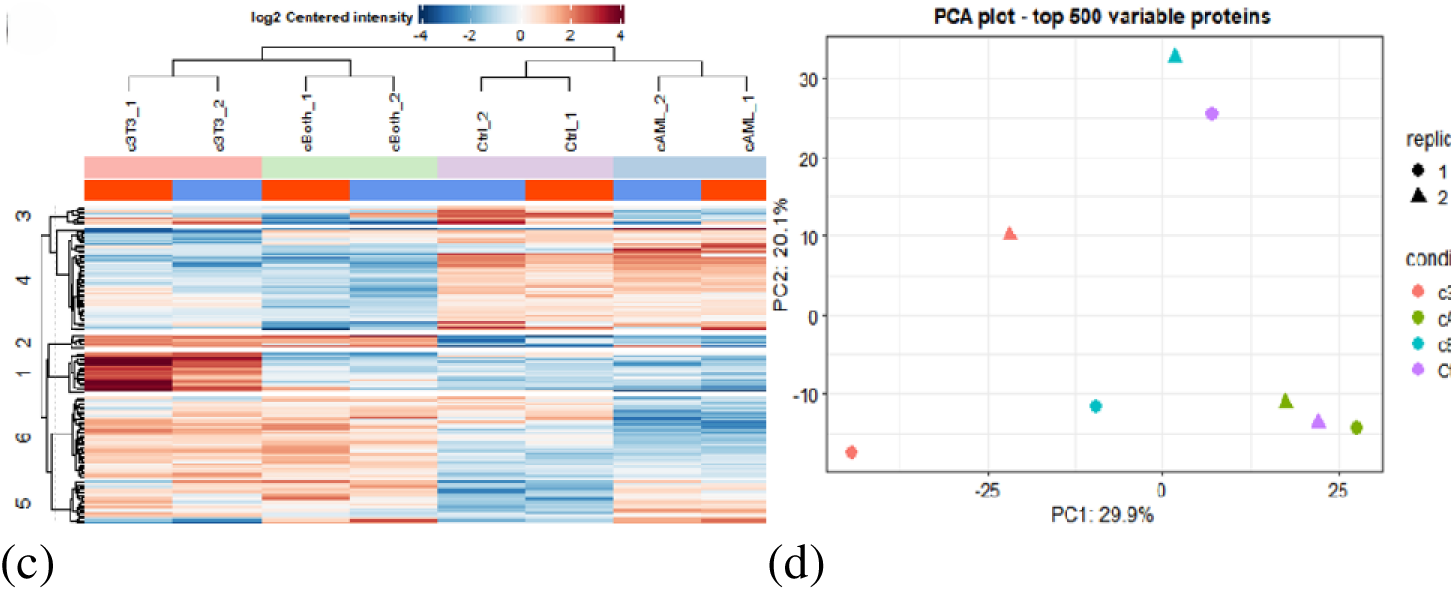
Heatmap showing hierarchical clustering of different treatment groups under different growth conditions. PCA plots are indicating separation between the different components; (**a**) Heatmap 2D (**b**) PCA plot 2D (**c**) Heatmap 3D (**d**) PCA plot 3D.

**Figure 3:**
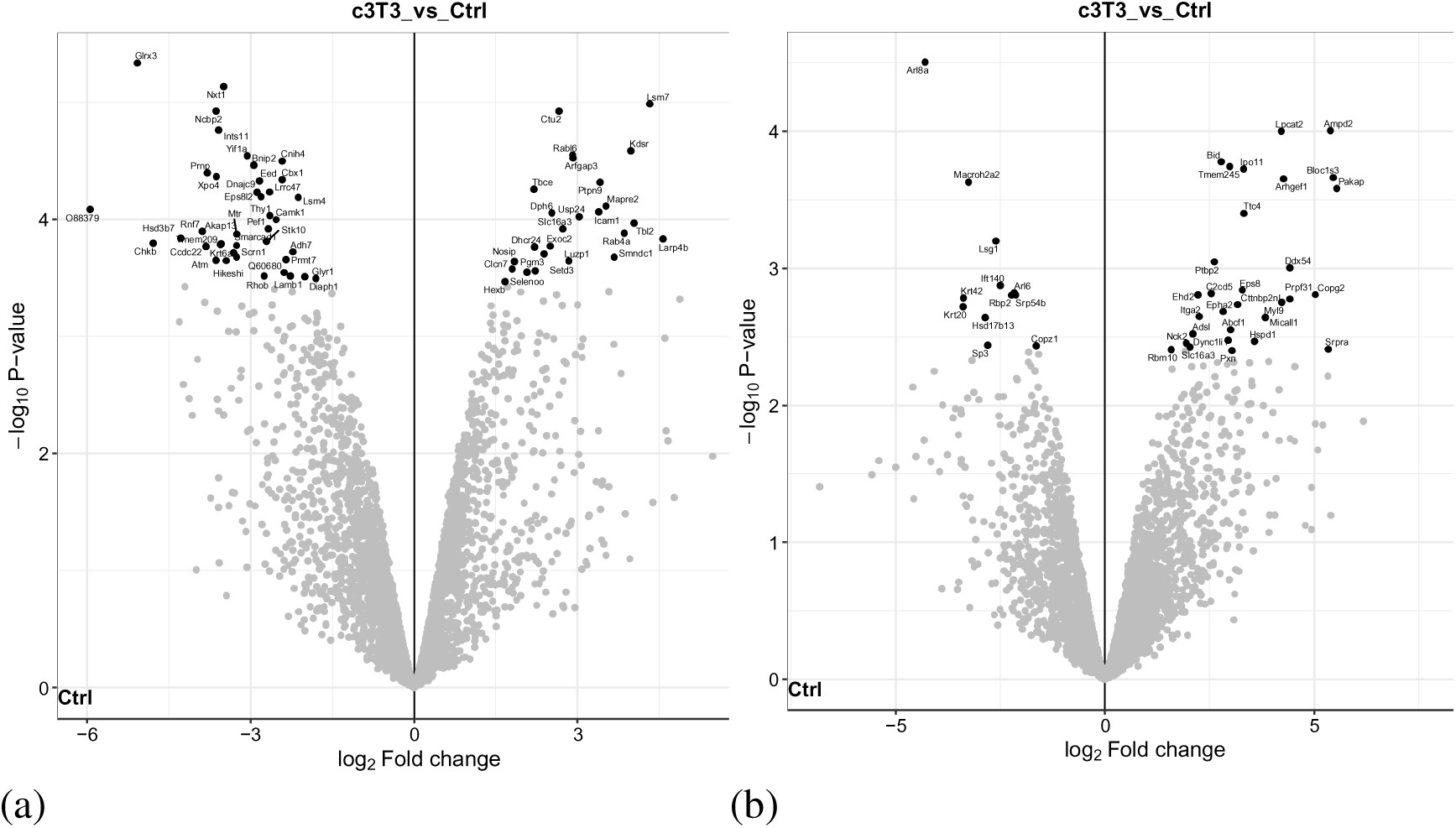
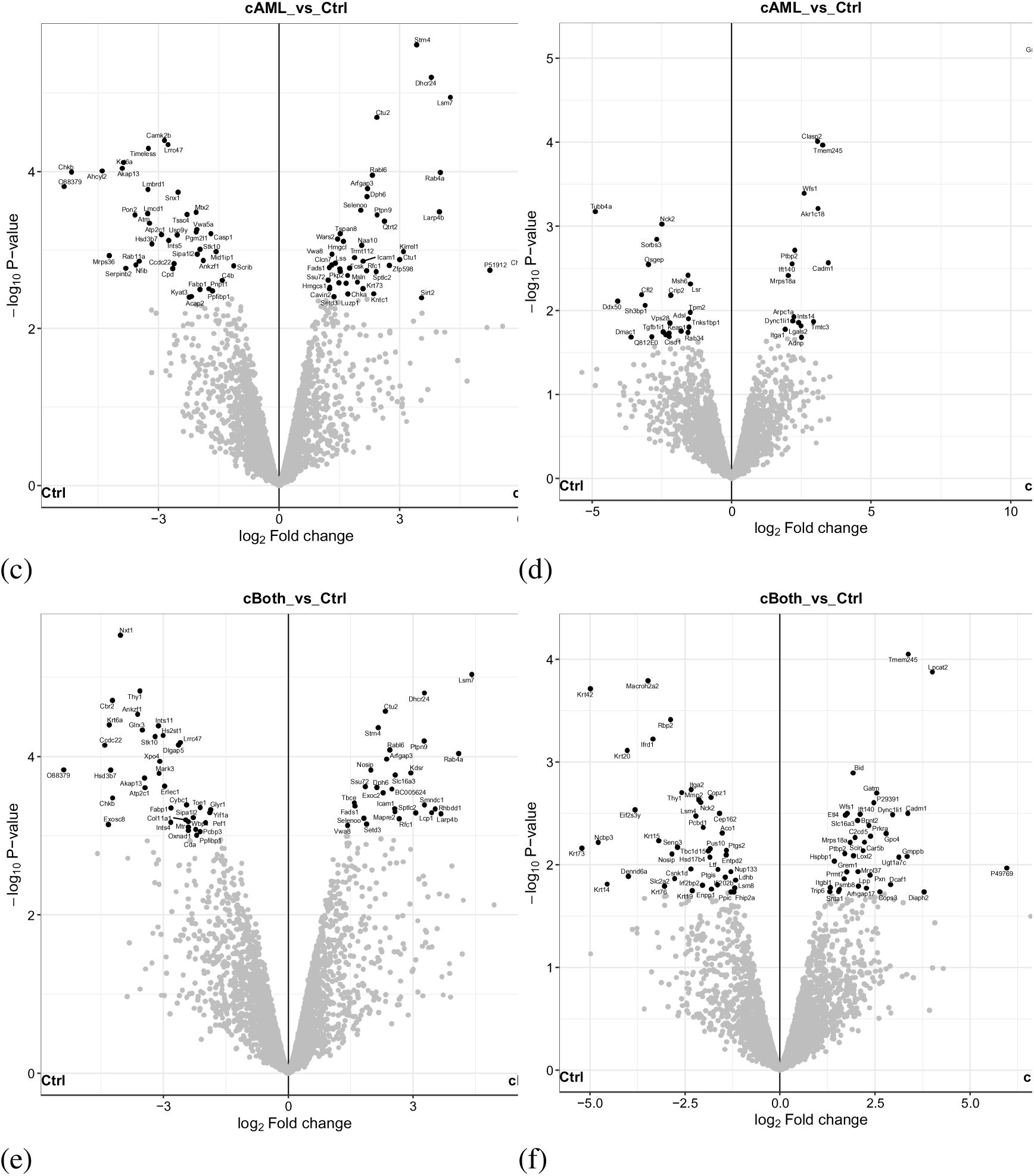
Volcano plots indicating significant differences in proteins between experimental groups identified using a two-sided t-test allowing for an FDR of 0.05 and mean fold change of log2(x) ± 1.0 for; (**a**) 3T3 versus ctrl 2D; (**b**) 3T3 versus ctrl 3D; (**c**) AML 12 versus ctrl 2D; (**d**) AML 12 versus ctrl 3D; (**e**) cBoth versus Ctrl 2D; (**f**) cBoth versus Ctrl 3D (Figure generated using RStudio software loaded with the DEP package.

### 3.2. DEP’s Identified in 3D Model showed Marked Enrichment in HCC associated Protein Classes

Functional enrichment analysis (Gene Ontology) of differentially expressed proteins was carried out to identify sets of pathways relevant with the HCC phenotype. GO ontology using PANTHER revealed that majority of the differentially expressed proteins (DEPs) belonged to the classes of cytoskeletal, RNA metabolism and scaffold/adaptor proteins. Expression of these protein classes was found to be stronger in all the groups of 3D (c3T3 14%, cAML 32% and cBoth 17%) compared to 2D (c3T3 4.4%, cAML1.70%, cBoth 4.4%). Moreover, structural proteins were only expressed in mixed 3D cultures of AML12 and 3T3 which indicates closer homology of 3D systems to in vivo system. Within 3D, cAML showed the highest expression of these protein classes hence a stronger association with the HCC phenotype (Figure 4 a-f).

**Figure 4:**
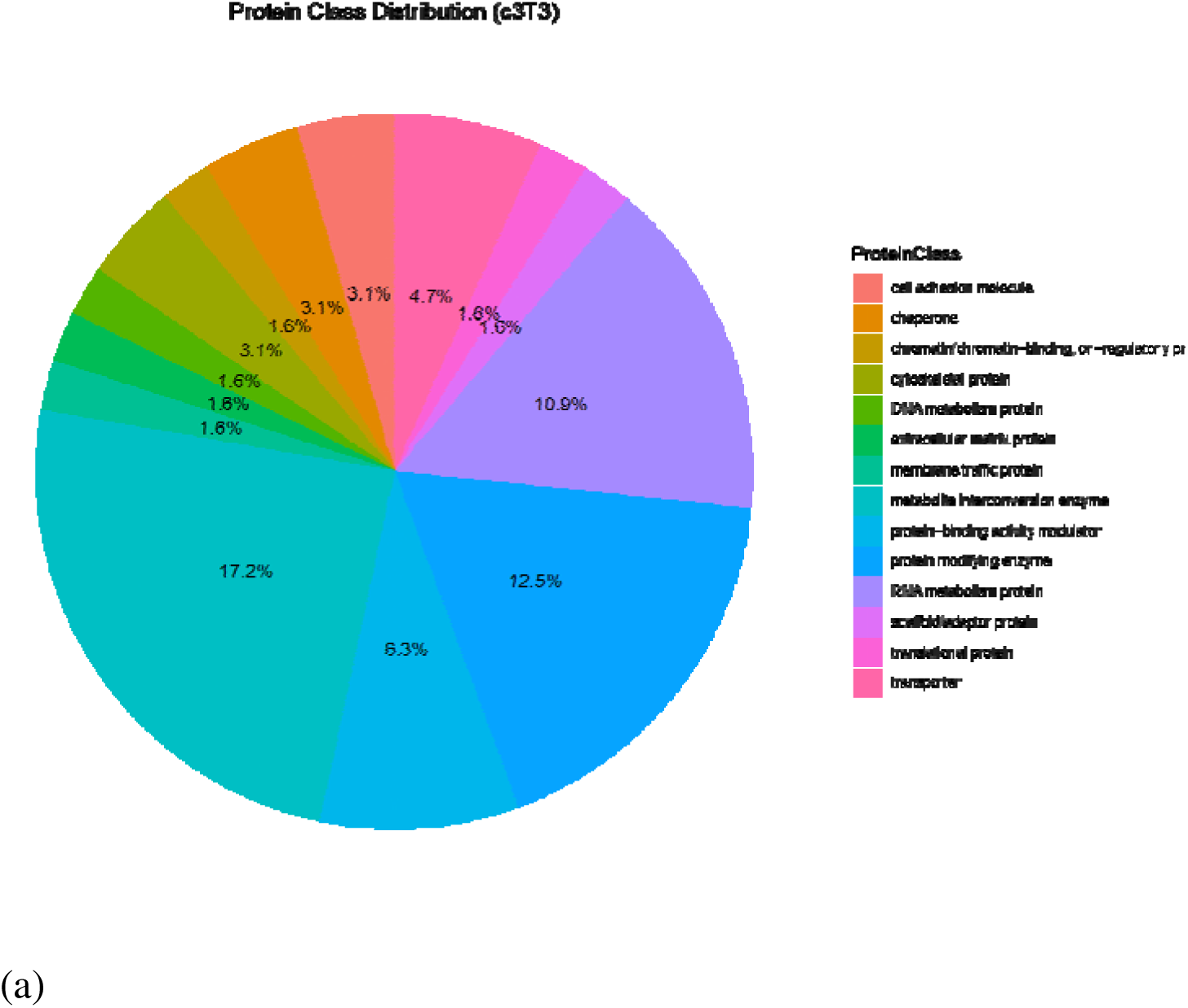
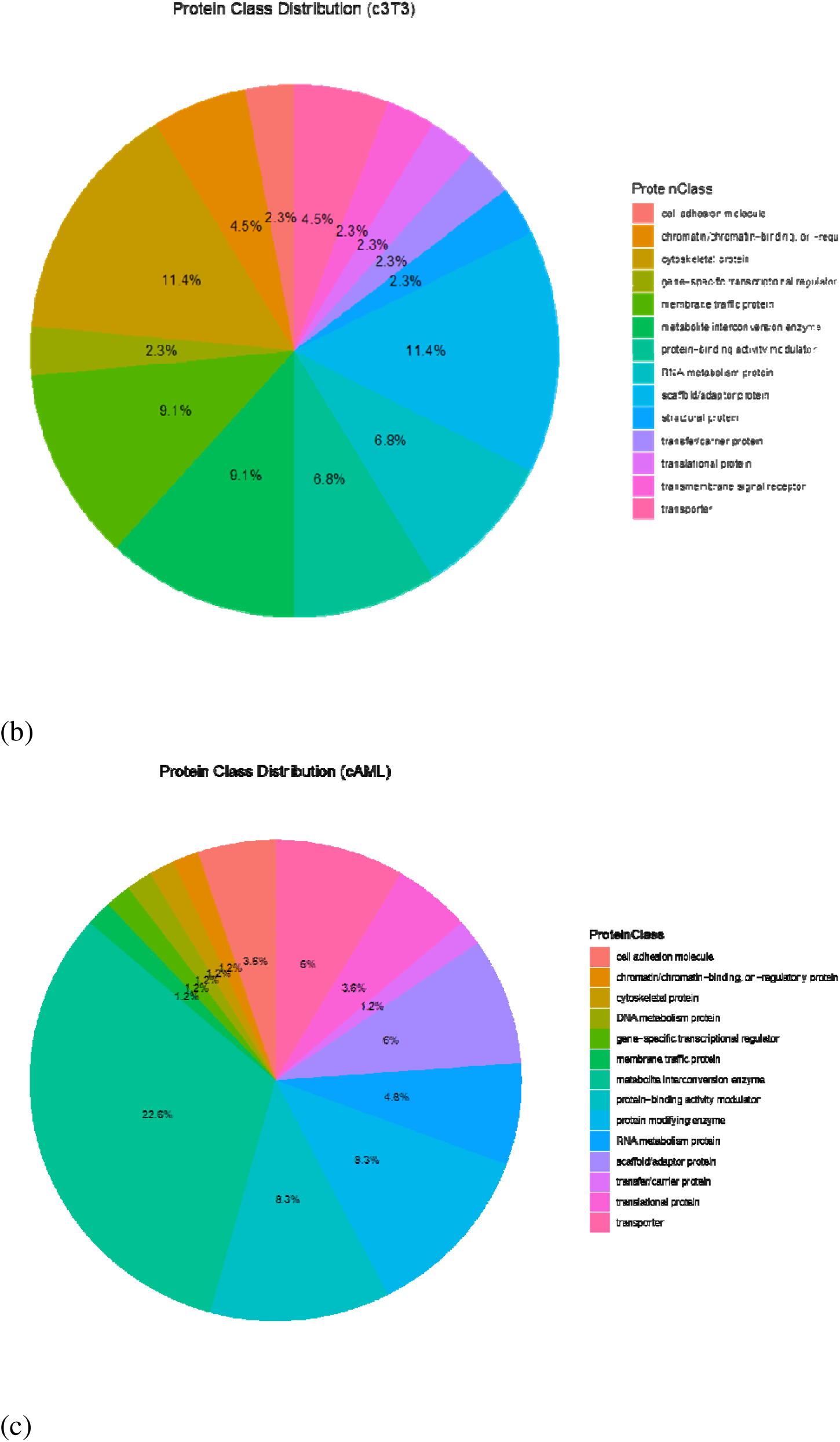
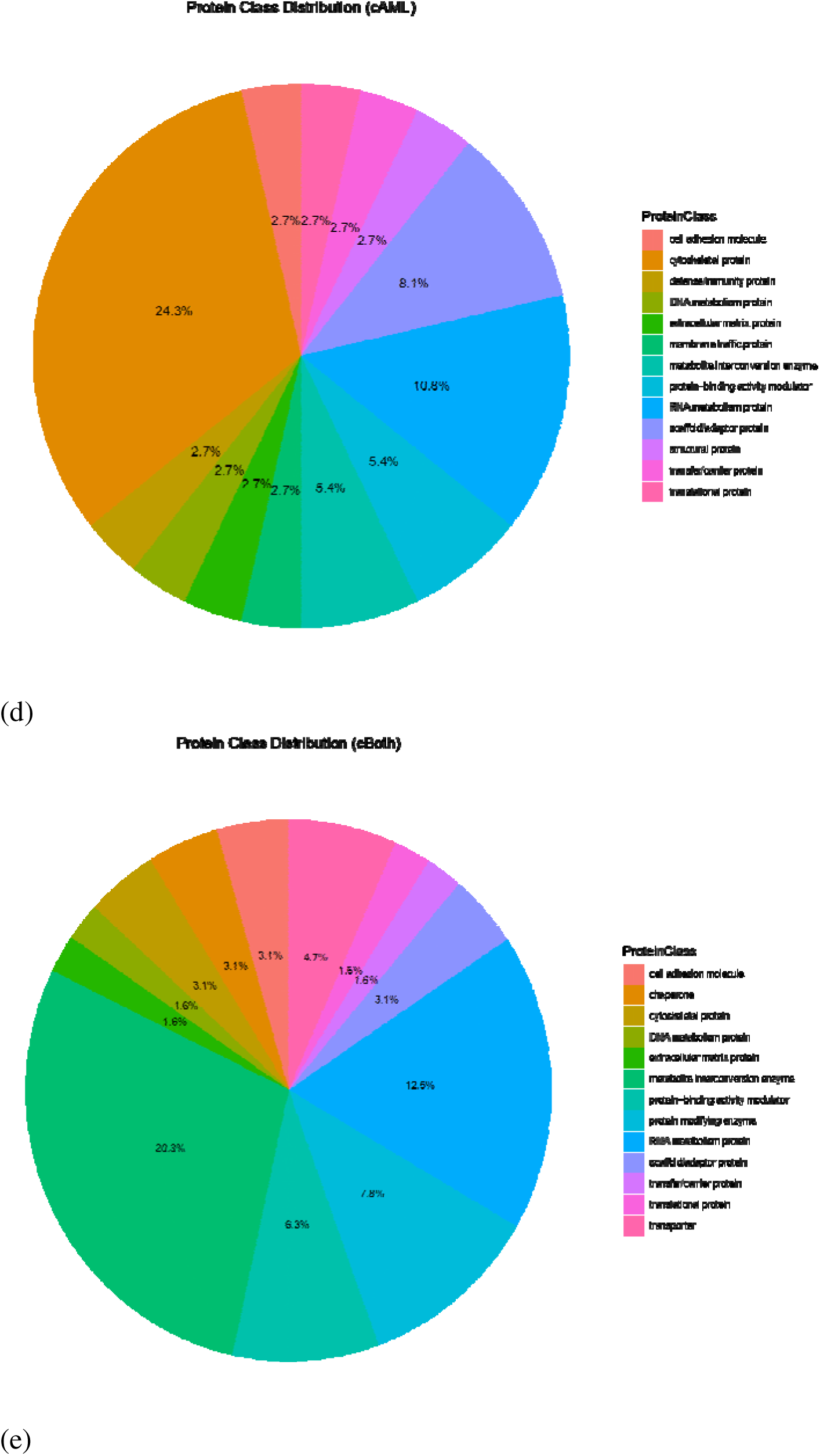
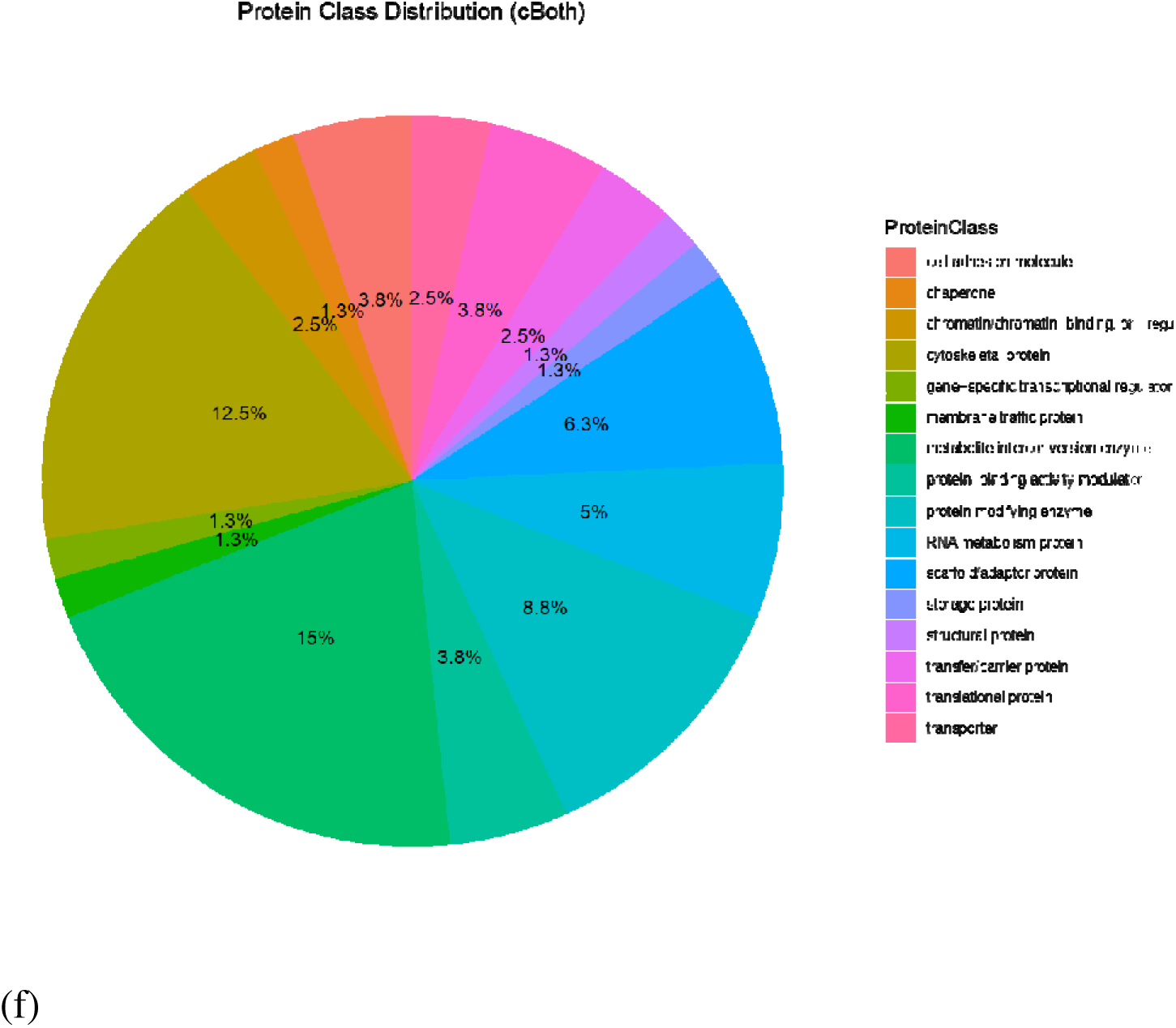
Pie charts generated through Panther represent protein classes. Each class is represented by a specific color code.; (**a**) 3T3 2D; (**b**) 3T3 3D; (**c**) cAML 2D; (**d**) cAML 3D; (**e**) cBoth 2D; (**f**) cBoth 3D.

### 3.3. Overrepresentation Analysis confirms HCC related pathway enrichment in 3D

GO analysis was performed in terms of c3T3, cAML and cBoth. In 2D, GO pathways were found to be associated with snRNA processing, tRNA metabolic process, postsynaptic endosome, nuclear-transcribed mRNA catabolic process and histone methyltransferase activity (Figure 5a). In 3D, GO pathways were associated with intermediate filament and cytoskeleton organization, actin filament organization, focal adhesion, cell-substrate junction, hydro-lyase and carbon-oxygen lyase activity, guanyl nucleotide and ribonucleotide binding, GTP binding, GTPase activity and protein localization (Figure 5b). GO analysis further validated that HCC related pathways were better enriched under 3D condition.

**Figure 5:**
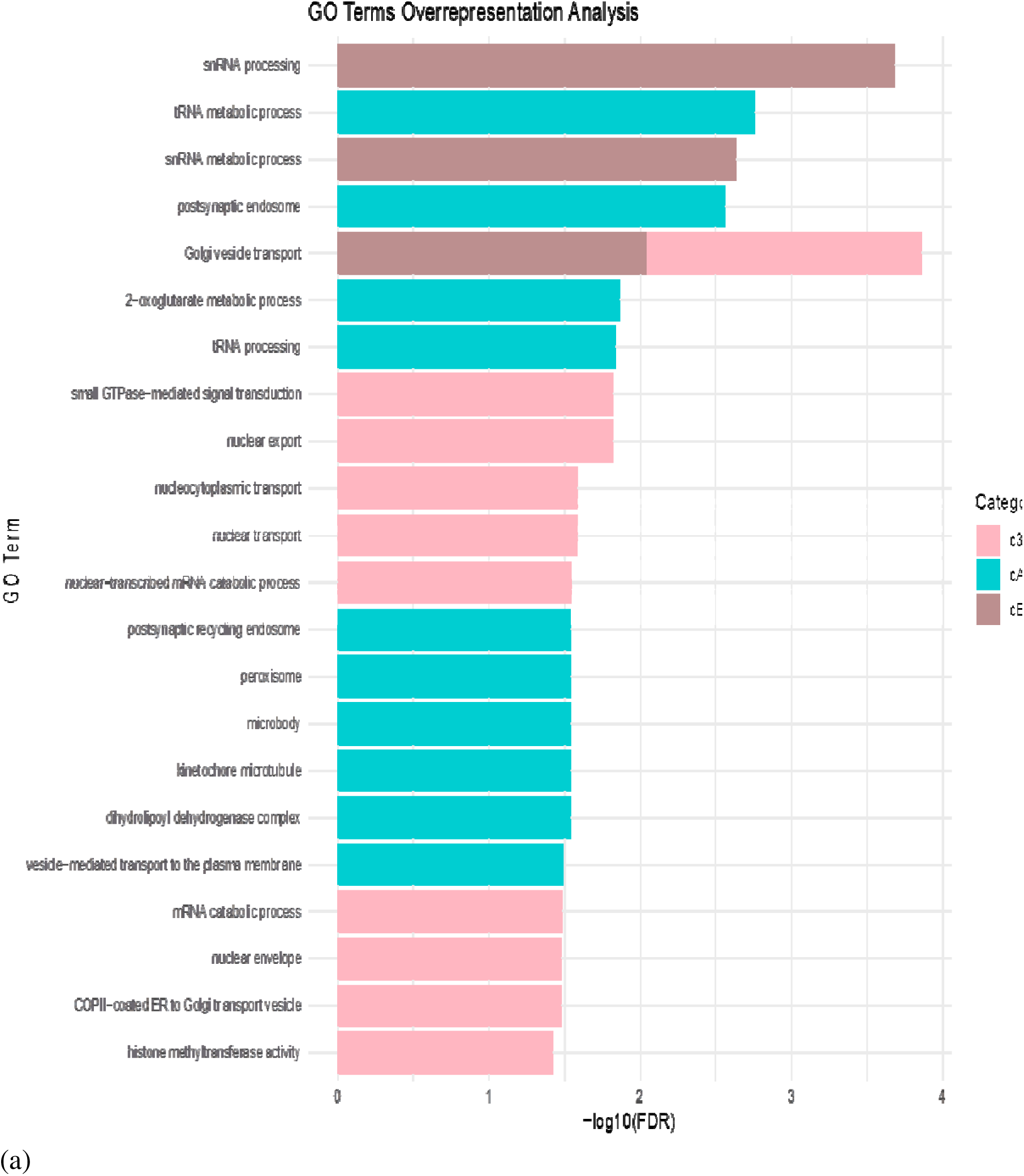
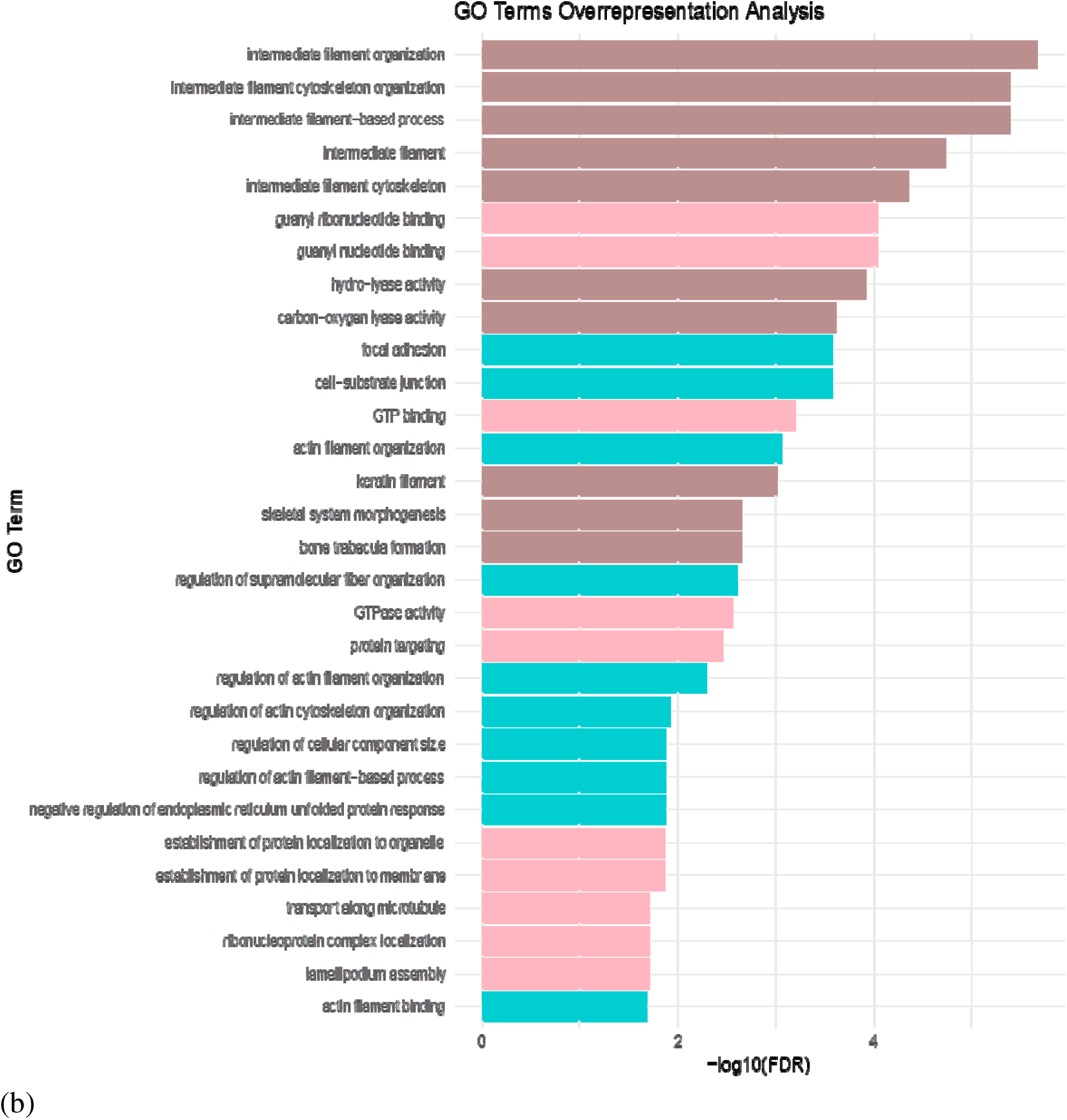
GO term analysis pink color indicates pathways enriched in 3T3 treated with DEN, green color indicates pathways enriched in AML12 treated with DEN and brown color indicates pathways enriched in the last group where both AML12 and 3T3 were treated with DEN; **(a)** GO term Analysis for 2D **(b)** GO term Analysis for 3D.

### 3.4. Molecular Complex Detection (MCODE) Cluster Module Identified Keratin Cluster in DEN Treated stromal-epithelial (cBoth) 3D model

Protein-protein interactions were generated by STRING with a medium confidence of 0.400. cboth showed highest MCODE score of 4.0 with 78 nodes and 35 edges in 3D culture (Krt 14, Krt 15, Krt 19, Krt 20). cAML and c3T3 shared the same MCODE score of 3.0. cAML showed 37 nodes and 11 edges while c3T3 showed 42 nodes and 14 edges (Figure 6 a-f). From here onwards only 3D cultures with three different experimental groups were selected for further analysis since 2D cultures didn’t show promising results.

**Figure 6:**
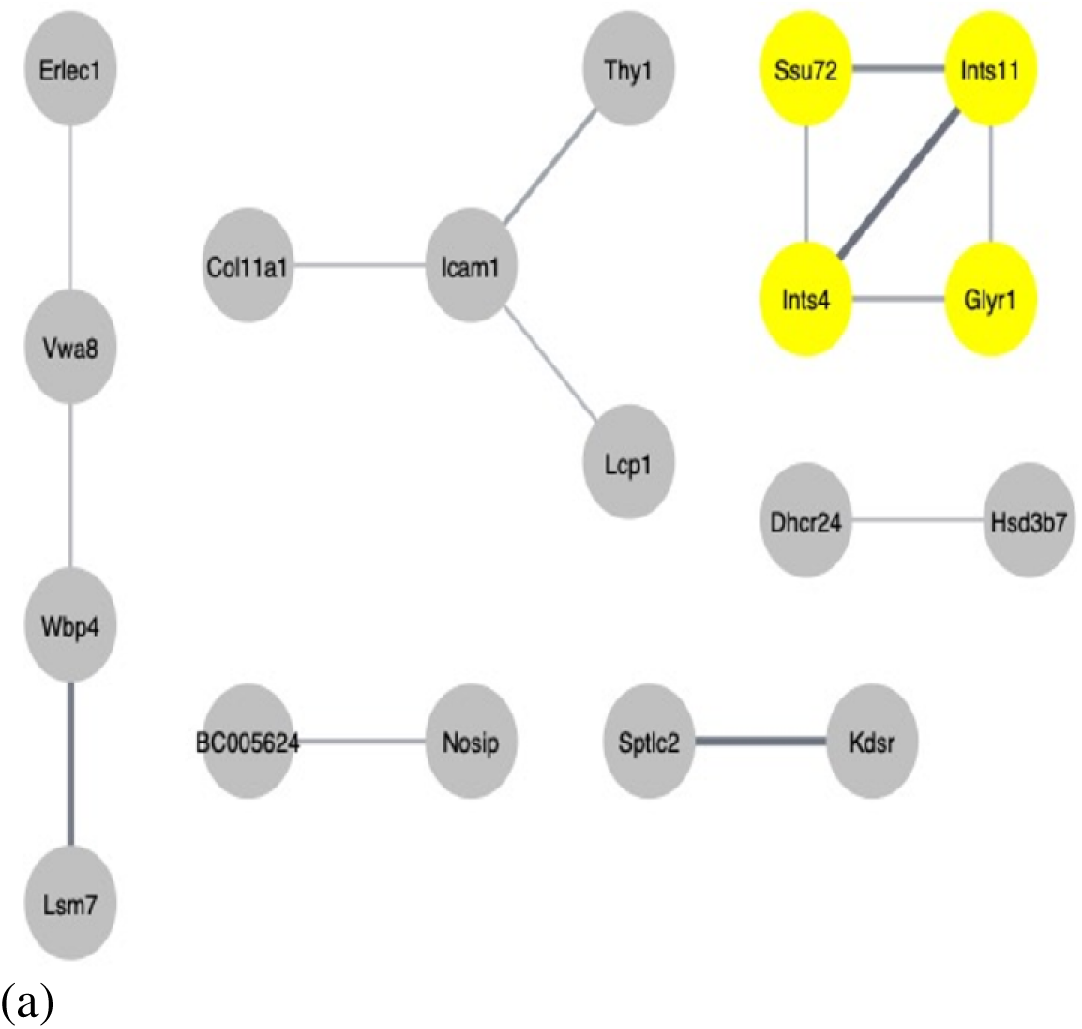
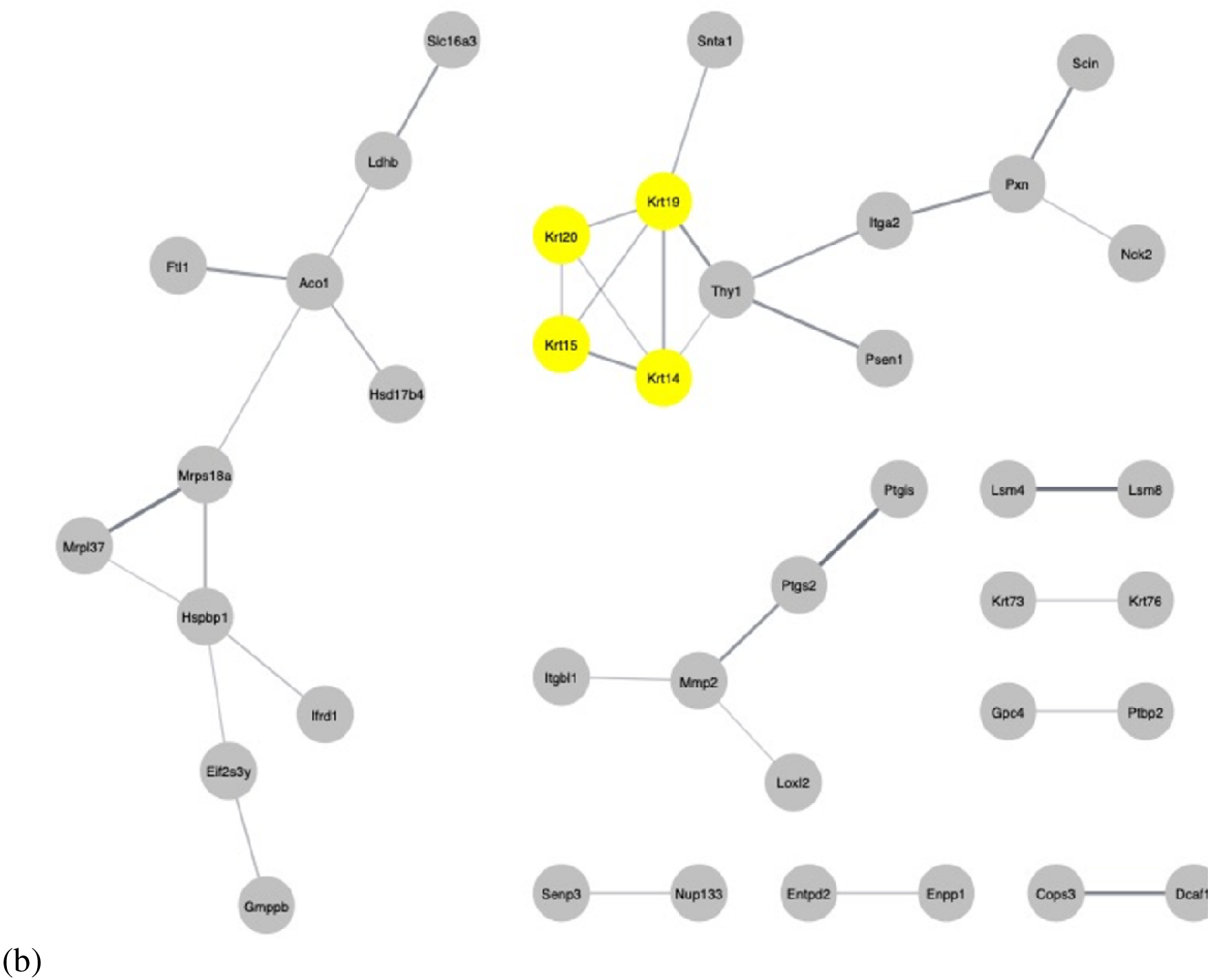
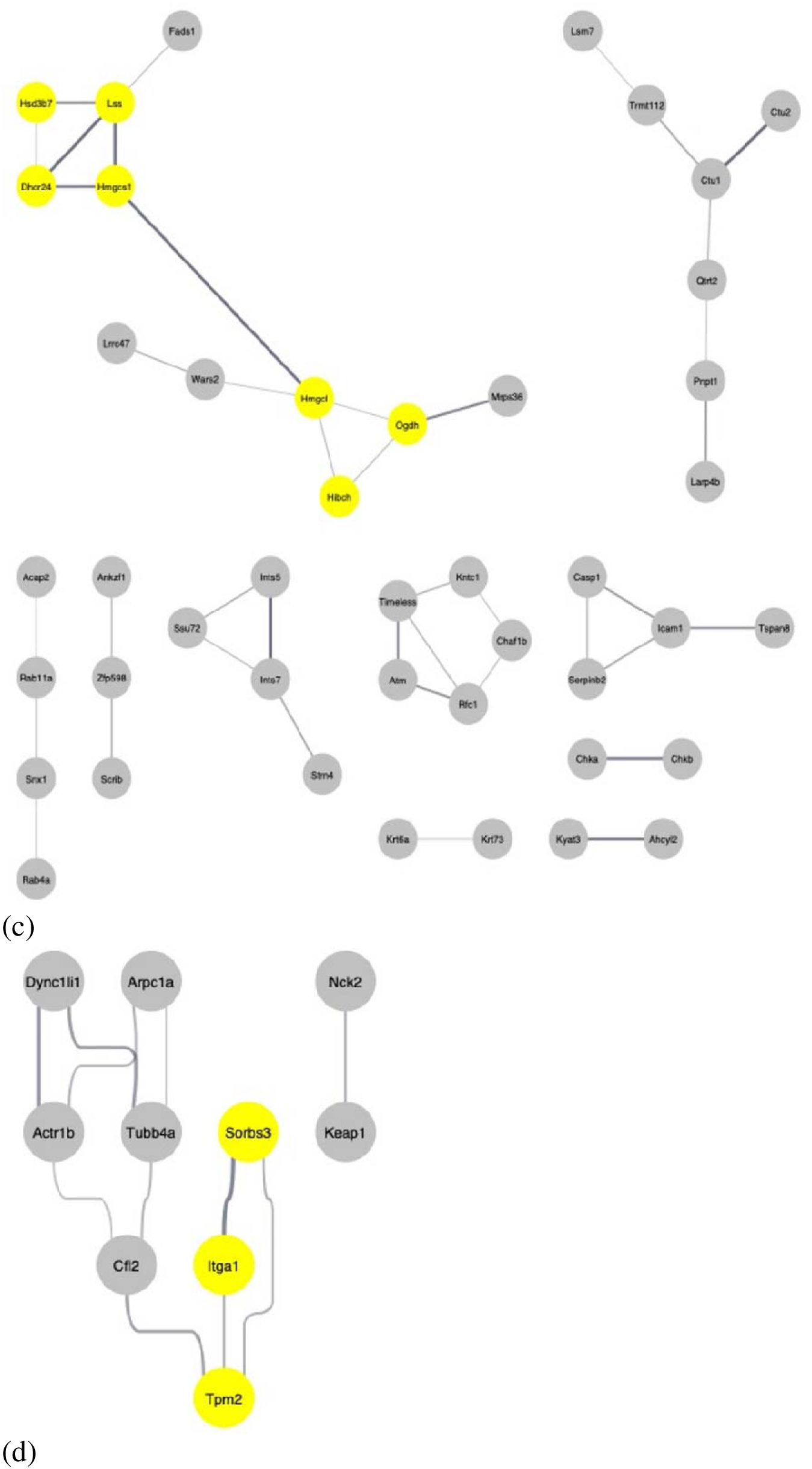
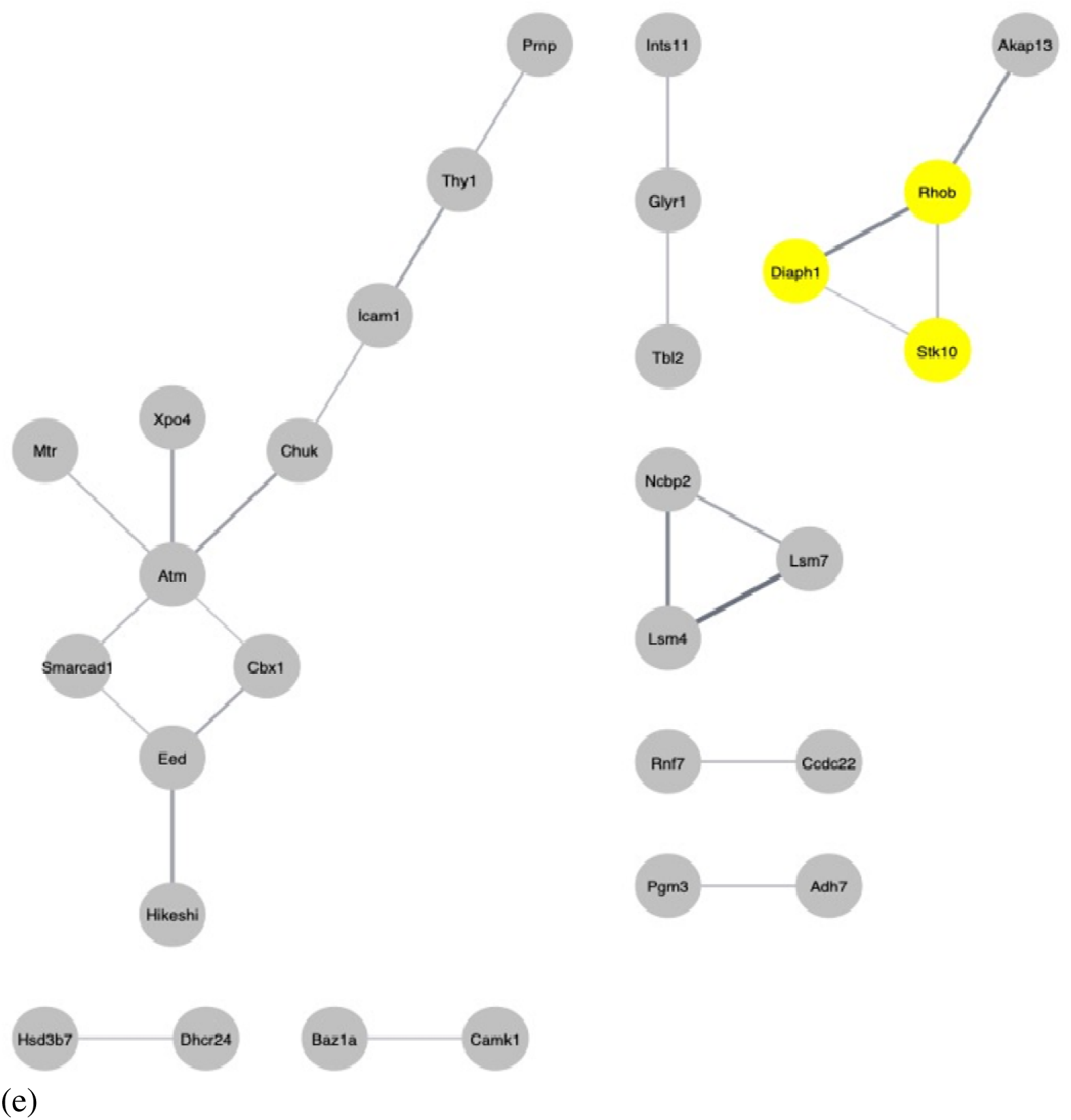
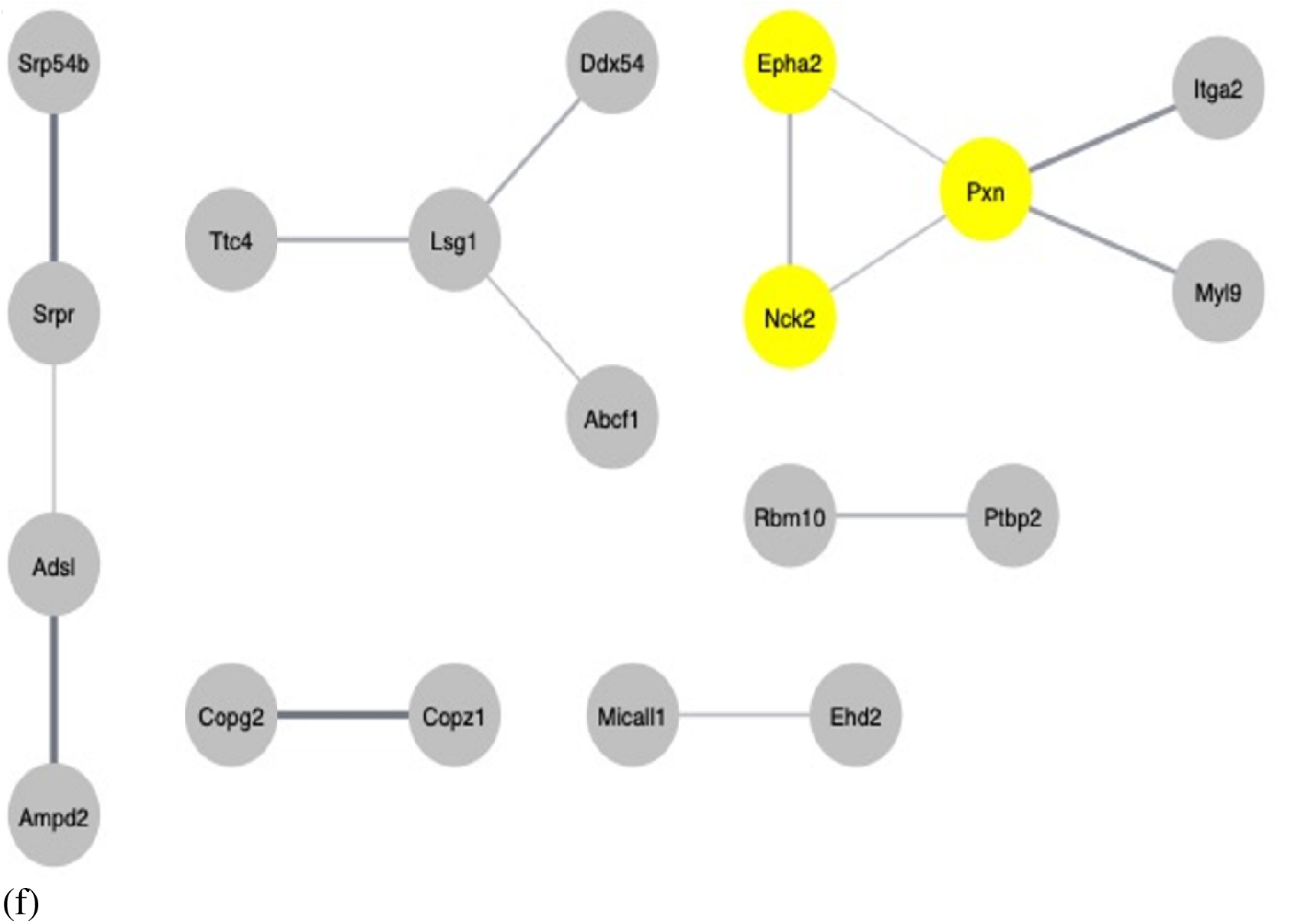
Protein-protein interactions (**a**) cBoth 2D; (**b**) cBoth 3D; (**c**) cAML_2D; (**d**) cAML_3D; (**e**) c3T3_2D; (**f**) c3T3_3D

### 3.5. Dysregulation of Keratin 14, keratin 15 and keratin 20 were Significantly Associated with Poor Patient Survival

The survival analysis of genes presents in the top cluster of PPI network indicated that only 1 protein PXN (P < 0.0099) from c3T3 was associated with poor overall survival. Three genes KRT14 (P< 0.026), KRT 15 (P < 0.022), KRT 20 (P<0.0012) from cBoth showed overall poor survival in HCC patients (Figure 7 a-d).

**Figure 7:**
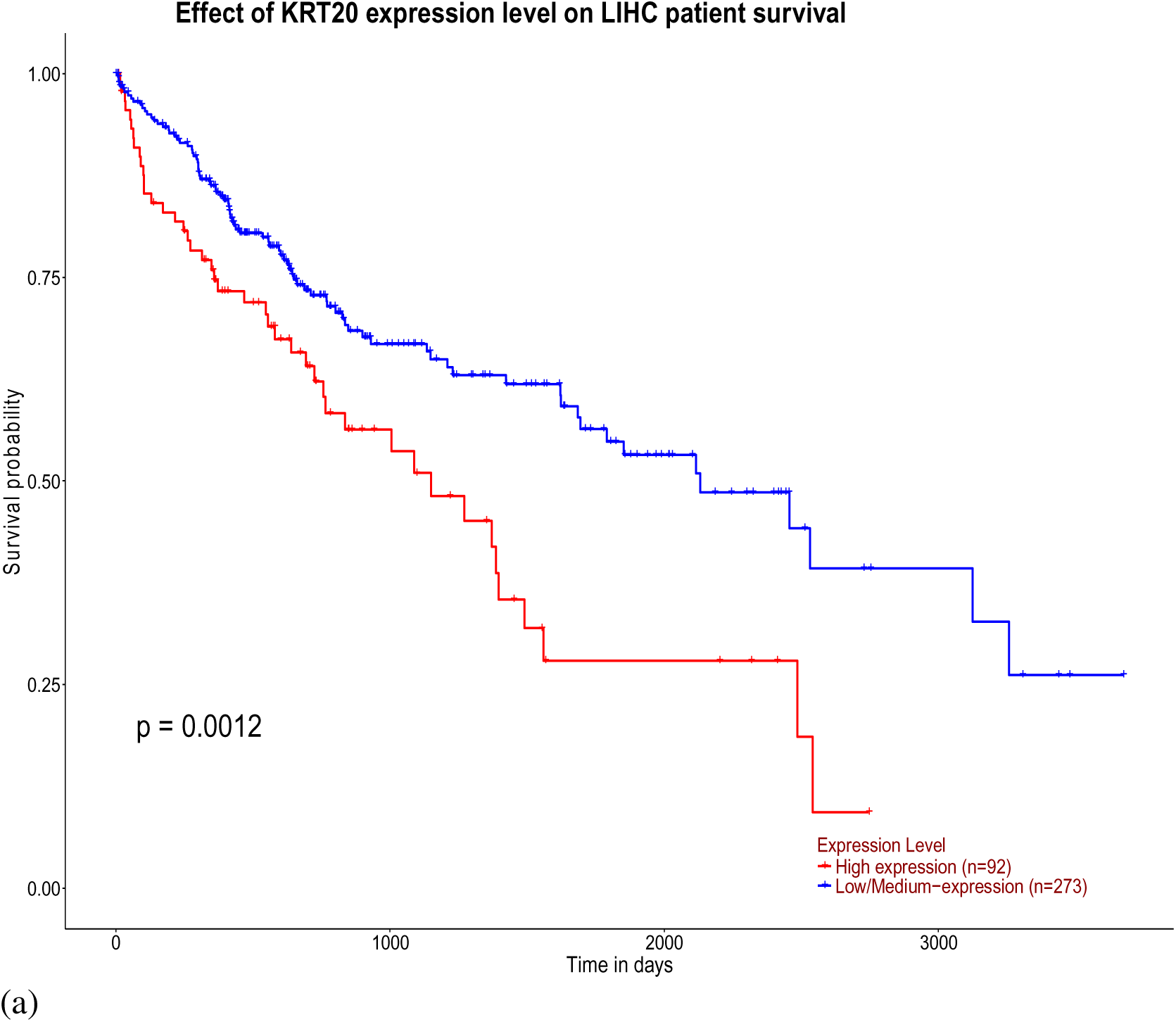
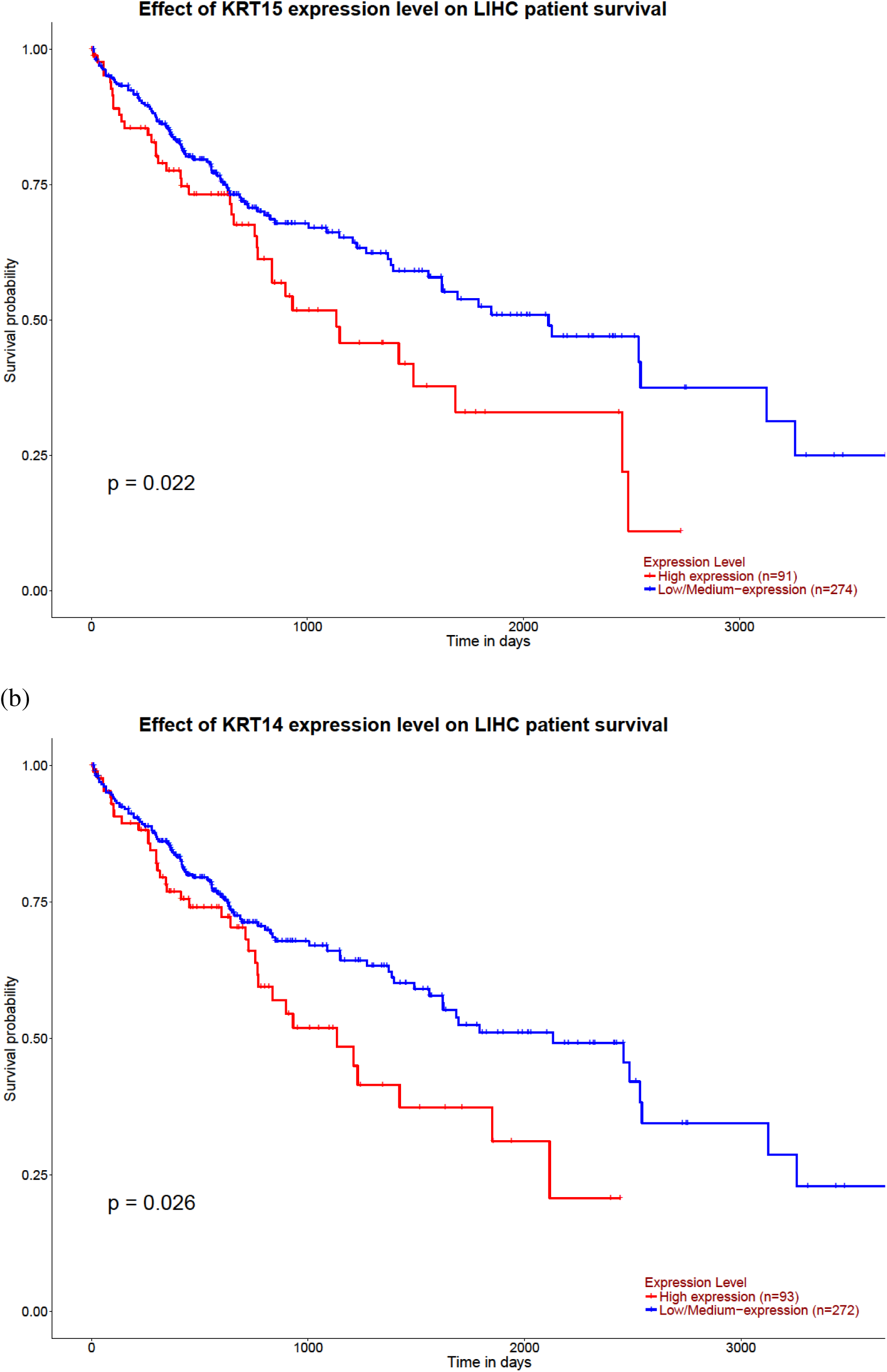
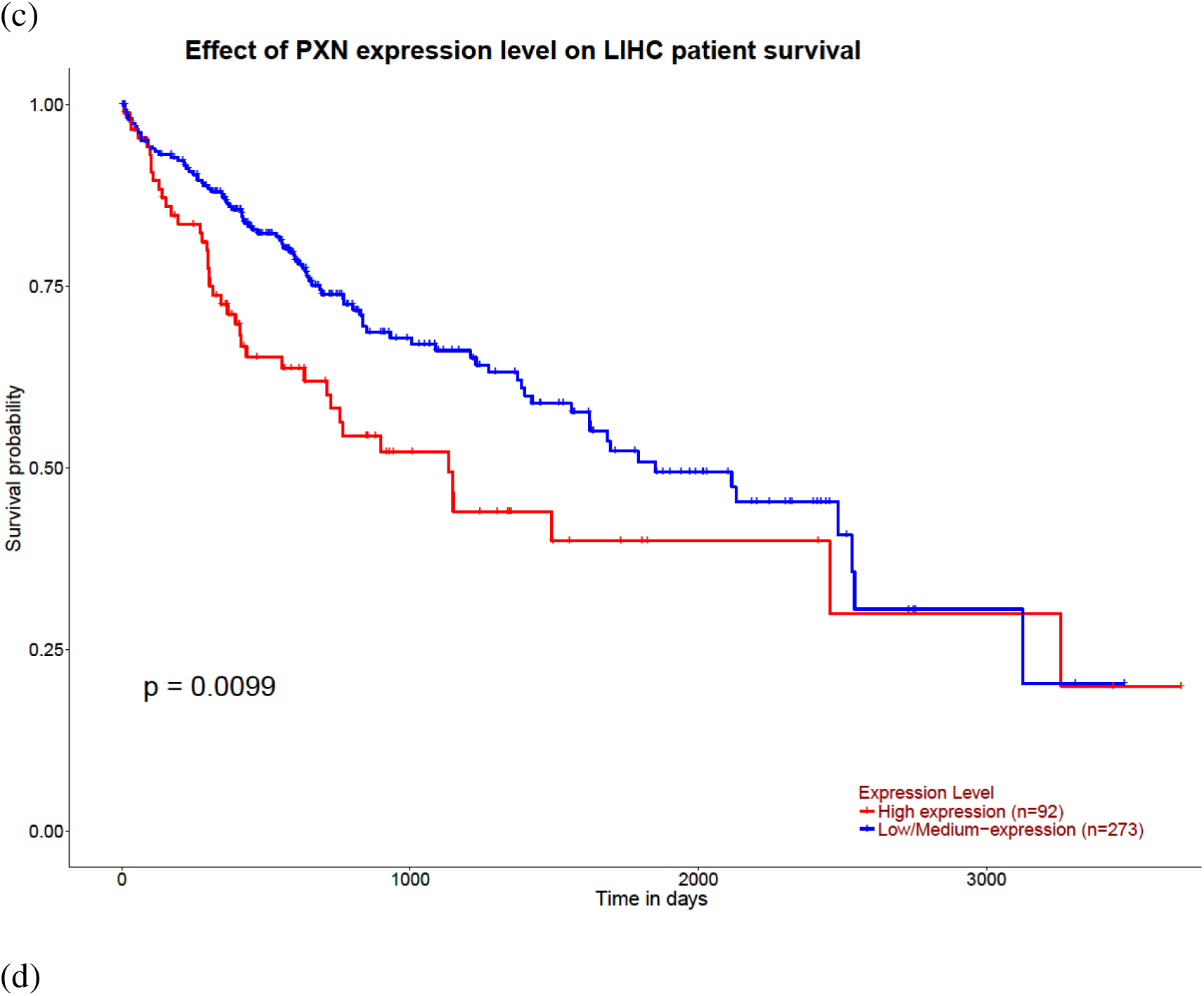
Survival analysis was carried out using UCLAN; (**a**) KRT20; (**b**) KRT 15; (**c**) KRT14; (**d**) PXN.

### 3.6. Dysregulated Keratin 8 and 18 show Maximum Divergence from Hepatocyte like Character

Expression of proteins encoding normal hepatocyte character was obtained from another study [15] was investigated in proteins expressed in 3D cultures. Proteins expressed in cultures treated with carcinogen were compared to control group to observe the expression of aforementioned proteins across different treatment groups. The heatmap was plotted on the basis of logFC values (Figure 8). Seventeen out of Forty-two proteins were represented in this data. In cAML six proteins namely, APOB (Apolipoprotein B), MAOA (Monoamine oxidase A), MAT2A (Methionine Adenosyltransferase 2A), ITIH3 (Inter-alpha-trypsin inhibitor heavy chain 2), KRT8 (Keratin 8) and KRT18 (Keratin 18) showed maximum divergence compared to the control sample in cAML. In cBoth four proteins namely, CFB (Complement factor B), ALB (Albumin), EPHX2 (Epoxide hydrolase 2) and SERPINC1(Serpin Family C member 1) showed maximum divergence. In 3T3 seven proteins showed maximum divergence in expression namely: EPHX1 (Epoxide hydrolase 1), PROX1 (Prospero homeobox 1), MAOB (Monoamine oxidase B), CPS1(Carbomoyl phosphate synthetase), ITIH2 (Inter-alpha-trypsin inhibitor heavy chain), AMBP (Alpha-1-microglobulin percursor), MAT1A (Methionine Adenosyltransferase 1A).

**Figure 8:**
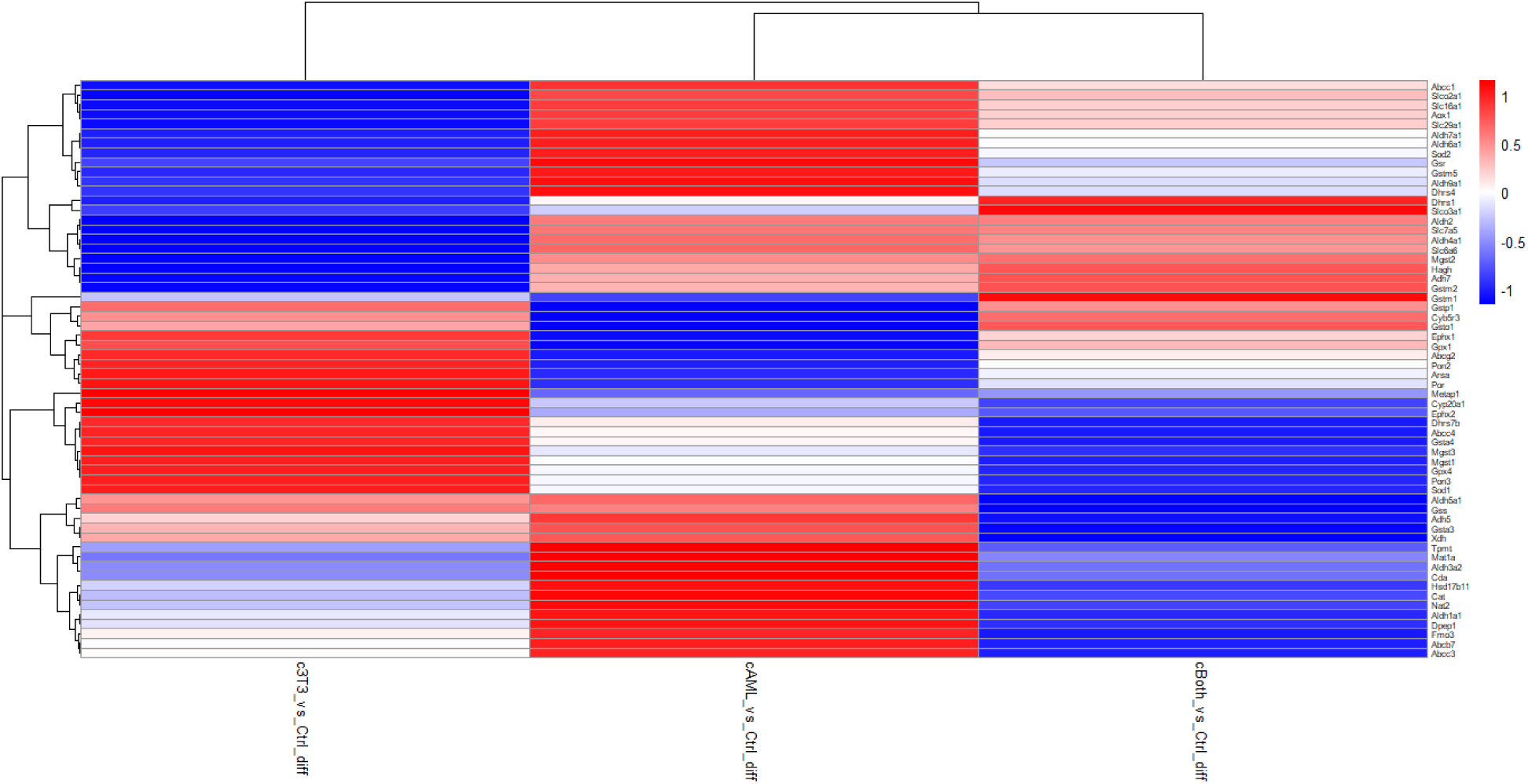
Heatmaps showing divergence from Hepatocyte like character

### 3.7. Validation of Bioinformatics Analysis through Immunohistochemical Assays

Immunocytochemical analysis of all DEN treated groups was carried out for tumor markers (collagen IV, PCNA, keratin 18 and E-Cadherin). Results indicated that the expression of collagen IV, PCNA and keratin 18 was lowest in An3n (both AML12 and 3T3 were treated with vehicle), whereas highest expression was observed in cBoth (both AML12 and 3T3 were treated with DEN. c3T3 and cAML showed intermediate levels of expression. E-Cadherin showed the reverse pattern of expression consistent with the HCC phenotype. The untreated samples showed highest level of E-Cadherin with a gradual decrease in expression in cAML and c3T3, while the least expression was observed in cBoth (Figure 9).

**Figure 9:**
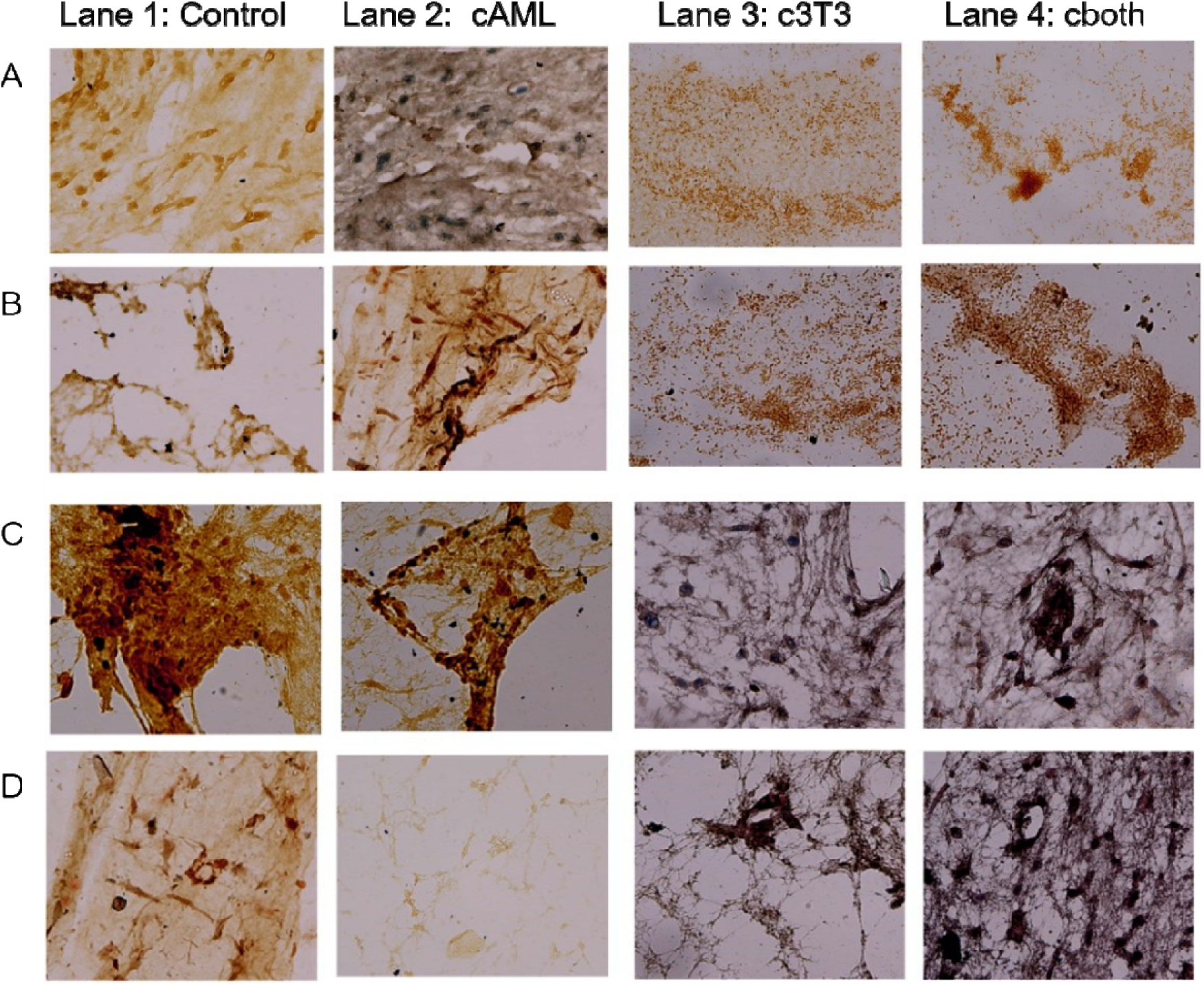
Left to right Lane 1: Control; Lane 2: cAML; Lane 3: c3T3; Lane 4: cBoth; (Row: A) (PCNA) (Row:B) (Keratin 18) (Row:C) E-Cadherin (Row:D) Collagen IV

## 4. Discussion

For the last one century cancer has been understood to have arisen from mutations in epithelial cells. The role of stroma in tumorigenesis and pathogenesis of cancer has long been ignored. It is quite recent that stroma has been demonstrated to play a significant role in the occurrence and progression of cancer through its intrinsic and acquired tumor related properties [16]. The purpose of this study was to determine which culture condition (2D or 3D) is best suited to study the initiation of HCC and whether the target of Diethylnitrosamine is liver epithelium or stroma or both stroma and epithelium. For the fulfilment of these goals both stromal and epithelial cell lines were exposed to DEN separately and then recombined into four different combinations. All four recombinants were cultured under both 2D and 3D culture conditions. Each recombinant cultured in 2D was compared to the corresponding recombinant in 3D culture to determine which of the two culture types offers better niche for supporting a more *in vivo* like phenotype. Under both culture conditions, all the recombinants (each containing DEN treated, stoma or epithelium or both) were compared among themselves to determine which tissue compartment is the target of DEN. The results obtained are expected to provide new insights into the proteomic permutations in liver caused by the exposure to DEN and the role of stroma in modulating these proteomic variations.

The proteomic landscape of 3D cultures demonstrated distinct differences compared to 2D cultures, reflecting their enhanced physiological relevance. Protein class distribution analysis using PANTHER revealed that structural proteins, particularly those involved in extracellular matrix organization, were significantly enriched in 3D hydrogel cultures across all three recombinant groups. Protein classes closely associated with HCC such as cytoskeletal proteins, RNA metabolism proteins and scaffold/adaptor proteins also showed stronger expression in 3D cultures (c3T3 14%, cAML 32% and cBoth 17%) compared to 2D culture (c3T3 4.4%, cAML1.70%, cBoth 4.4%) indicating that 3D culture supports stronger protein expression patterns. The advantage of 3D culture system over 2D was also consistently reflected in GO term analysis and protein-protein interactions using Cytoscape. Our findings are consistent with the findings of another group [15] who have shown before that global proteomic analysis of primary human hepatocytes in 3D cultures displayed greater temporal stability but higher sensitivity to hepatic toxins as opposed to the 2D sandwich cultures that showed greater proteomic alteration over the period of 14 days and lesser sensitivity to hepatotoxins. Hence 3D culture of liver cells reflects greater metabolic capacity and competence compared to the 2D cultures [17].

MCODE revealed that Keratins (KRT 14, KRT 15, KRT 19 and KRT 20) exhibited the largest network in DEPs of cBoth where both epithelial and stromal cells were exposed to DEN. Kaplan-Meier (KM) analysis of cBoth further indicated that all the proteins (KRT 14, KRT 15 and KRT 20) in the cluster were associated with poor patient survival. These findings are consistent with the literature that shows that various keratins are associated with different stages of HCC. K19 is a marker of tumor invasion and poor prognosis in HCC [18]. K15 levels have been found to be upregulated in most liver cancer cell lines [19]. KRT14 is also over expressed in various cancers including melanoma of [20].

Keratins are the intermediate filament forming protein family, closely associated with occurrence and progression of various cancers [20]. Keratin 8 and keratin 18 are the intermediate filament proteins expressed in hepatocytes which act to protect the cell from mechanical stress and provide structural stability. They have also been found to potentially increase the risk of HCC incidence [21]. In our study, heatmap of divergence from hepatocyte like character indicated that KRT18 was also downregulated. It has also been previously reported [22] that KRT18 interacts with several structural proteins in hepatocellular carcinoma including plectin, which is a multifunctional cytoplasmic cross-linked protein with multiple binding sites for cytoskeletal components. It also serves to connect different cytoskeletal components to form a complete and stable cytoskeletal network. In HCC, downregulated KRT18 contributes to cytoskeletal disturbances, nuclear instability and vulnerability. Another study [23] reported that the crosstalk between malignant epithelial cells and CAFs regulates the expression of keratin 19 in biologically aggressive HCC, suggesting the role of keratins in stromal epithelial interactions during the development of HCC. The role of keratins in liver cancer, particularly HCC has not been extensively studied so far [19], possibly because most cancers including HCC have been viewed to arise from epithelial cells. As the recent studies are unveiling the role of stroma in cancer, role of keratins in HCC is also becoming more and more evident. To the best of our knowledge this is the 1^st^ report indicating the potential role of keratin 14, 15 and 20 in relation to HCC.

Moreover, no protein from cAML was found to be associated with poor survival whereas only 1 protein paxillin (PXN) from c3T3 was found to be associated with poor patient outcomes. It has also been reported before [24] that strong upregulation of PXN participates in ITGB1 regulated cell cycle progression in HCC. PXN-AS1-L has also been identified to play oncogenic roles in HCC [25]. These findings suggest that the DEN targets both epithelium and stroma and the cross talk between epithelium and stroma is responsible for the initiation of HCC. The results of ICCs also validate these findings.

## 5. Conclusions

In conclusion, cancers are not only composed of cancer cells alone rather they are complex ‘ecosystems’ consisting of multiple cell types and noncellular factors. Our study highlights the power of 3D co-cultures of liver epithelial and stromal cells to replicate a biomimetic liver model, providing a more accurate representation of the *in vivo* tumor microenvironment. Stroma and epithelium when treated together with the carcinogen instead of either one of them getting the carcinogen, give rise to disruption of the keratin proteins (14,15,18, 20) whose human homologs are associated with poor survival. This underscores the translational relevance of our model, which mimics key features of human liver cancer.

This further leads us to hypothesize that since stromal cells act a mechanical barrier to the initiation of carcinogenesis by curtailing the aberrant cell proliferation, DEN exposure disrupts this balance by simultaneously targeting both liver stroma and epithelium. This causes the epithelium to break the mechanical barrier of stroma and recruit them as CAF and at the same time CAF acts on liver epithelium to contribute to liver cancer progression. The interplay between these compartments highlights the importance of stromal–epithelial interactions in carcinogenesis, reinforcing the potential of this 3D co-culture model as a platform for studying early tumorigenesis and testing therapeutic interventions in liver cancer.

## Author Contributions

Conceptualization, Salmma Salamah Salihah. and Sana Mahmood; methodology, Salma Salamah Salihah; Software, Bareera Bibi, Agnese Rizzoli; validation, Bareera and Salmma Salamah Salihah; resources, Asma Gul, Simoni Sidoli, Muhammad Tahir; data curation, Agnese Rizzoli, Salmma Salamah Salihah; writing—original draft preparation, Salmma Salamah Salihah; writing—review and editing, Muhammad Tahir, Agnese Rizzoli, Sana Mahmood and Sehrish Khan, Massoud vosough; visualization, Bareera Bibi, Salmma Salamah Salihah and Sehrish Khan; supervision, Asma Gul, Simoni Sidoli; project administration, Asma Gul; funding acquisition, Simoni Sidoli and Asma Gul. All authors have read and agreed to the published version of the manuscript.

## Funding

This research received no external funding.

## Data Availability Statement

Proteomics raw data were deposited on the ProteomeXchange (PRIDE) database under the project accession PXD063112. All the relevant data supporting the findings of this study is included in the article and supplementary materials. Any additional information can be obtained upon reasonable request from the corresponding author.

## Acknowledgments

The Sidoli lab gratefully acknowledges for funding the Hevolution Foundation (AFAR), the Einstein-Mount Sinai Diabetes center, and the NIH Office of the Director (S10OD030286).

## Conflicts of Interest

The authors declare no conflicts of interest.

## Disclaimer/Publisher’s Note

The statements, opinions and data contained in all publications are solely those of the individual author(s) and contributor(s) and not of MDPI and/or the editor(s). MDPI and/or the editor(s) disclaim responsibility for any injury to people or property resulting from any ideas, methods, instructions or products referred to in the content. referred to in the content.

## Notes

https://www.ebi.ac.uk/pride/

### Competing Interest Statement

The authors have declared no competing interest.

## References

1. Jubelin, C.; Muñoz-Garcia, J.; Griscom, L.; Cochonneau, D.; Ollivier, E.; Heymann, M.F.; Vallette, F.M.; Oliver, L.; Heymann, D. Three-dimensional in vitro culture models in oncology research. Cell Biosci. 2022, 12, 155. 10.1186/s13578-022-00887-3

2. Habanjar, O.; Diab-Assaf, M.; Caldefie-Chezet, F.; Delort, L. 3D cell culture systems: Tumor application, advantages, and disadvantages. Int. J. Mol. Sci. 2021, 22, 12200. 10.3390/ijms222212200

3. Ellero, A.A.; van den Bout, I.; Vlok, M.; Cromarty, A.D.; Hurrell, T. Continual proteomic divergence of HepG2 cells as a consequence of long-term spheroid culture. Sci. Rep. 2021, 11, 10917. 10.1038/s41598-021-89907-9

4. Ma, Y.; Hu, L.; Tang, J.; Guo, W.; Feng, Y.; Liu, Y.; Tang, F. Three-dimensional cell co-culture liver models and their applications in pharmaceutical research. Int. J. Mol. Sci. 2023, 24, 6248. 10.3390/ijms24076248

5. Qi, W.; Zhang, Q. Insights on epithelial cells at the single-cell level in hepatocellular carcinoma prognosis and response to chemotherapy. Front. Pharmacol. 2023, 14, 1292831. 10.3389/fphar.2023.1292831

6. Heindryckx, F.; Gerwins, P. Targeting the tumor stroma in hepatocellular carcinoma. World J. Hepatol. 2015, 7, 165–176. 10.4254/wjh.v7.i2.165

7. Zhao, Y.; Shen, M.; Wu, L.; Yang, H.; Yao, Y.; Yang, Q.; Du, J.; Liu, L.; Li, Y.; Bai, Y. Stromal cells in the tumor microenvironment: accomplices of tumor progression? Cell Death Dis. 2023, 14, 587. 10.1038/s41419-023-06110-6

8. Wu, J.; Liang, C.; Chen, M.; Su, W. Association between tumor-stroma ratio and prognosis in solid tumor patients: a systematic review and meta-analysis. Oncotarget 2016, 7, 68954–68965. 10.18632/oncotarget.12135

9. Al Hrout, A.; Cervantes-Gracia, K.; Chahwan, R.; Amin, A. Modelling liver cancer microenvironment using a novel 3D culture system. Sci. Rep. 2022, 12, 8003. 10.1038/s41598-022-11641-7

10. Bradford, M.M. A rapid and sensitive method for the quantitation of microgram quantities of protein utilizing the principle of protein-dye binding. Anal. Biochem. 1976, 72, 248–254. 10.1016/0003-2697(76)90527-3

11. Ingelman-Sundberg, M.; Lauschke, V.M. 3D human liver spheroids for translational pharmacology and toxicology. Basic Clin. Pharmacol. Toxicol. 2022, 130 (Suppl. 1), 5–15. 10.1111/bcpt.13587

12. Gieseck, R.L., 3rd; Wilson, M.S.; Wynn, T.A. Type 2 immunity in tissue repair and fibrosis. Nat. Rev. Immunol. 2018, 18, 62–76. 10.1038/nri.2017.90

13. Bhatia, S.N.; Ingber, D.E. Microfluidic organs-on-chips. Nat. Biotechnol. 2014, 32, 760–772. 10.1038/nbt.2989

14. Maffini, M.V.; Soto, A.M.; Calabro, J.M.; Ucci, A.A.; Sonnenschein, C. The stroma as a crucial target in rat mammary gland carcinogenesis. J. Cell Sci. 2004, 117, 1495–1502. 10.1242/jcs.01000

15. Valkenburg, K.C.; de Groot, A.E.; Pienta, K.J. Targeting the tumour stroma to improve cancer therapy. Nat. Rev. Clin. Oncol. 2018, 15, 366–381. 10.1038/s41571-018-0007-1

16. 16. Ellero, A. A., van den Bout, I., Vlok, M., Cromarty, A. D., and Hurrell, T. (2021). Continual proteomic divergence of HepG2 cells as a consequence of long-term spheroid culture. Sci. Rep. 11(1), 10917. 10.1038/s41598-021-89907-9

17. Yang, S.; Ooka, M.; Margolis, R.J.; Xia, M. Liver three-dimensional cellular models for high-throughput chemical testing. Cell Rep. Methods 2023, 3, 100432. 10.1016/j.crmeth.2023.100432

18. Kawai, T.; Yasuchika, K.; Ishii, T.; Katayama, H.; Yoshitoshi, E.Y.; Ogiso, S.; Kita, S.; Yasuda, K.; Fukumitsu, K.; Mizumoto, M.; Hatano, E.; Uemoto, S. Keratin 19, a cancer stem cell marker in human hepatocellular carcinoma. Clin. Cancer Res. 2015, 21, 3081– 3091. 10.1158/1078-0432.CCR-14-1936

19. Wang, J.; Zhu, G. Silencing of keratin 15 impairs viability and mobility while facilitating the doxorubicin chemosensitivity by inactivating the β catenin pathway in liver cancer. Oncol. Lett. 2023, 26, 447. 10.3892/ol.2023.14034

20. Han, W.; Hu, C.; Fan, Z.J.; Shen, G.L. Transcript levels of keratin 1/5/6/14/15/16/17 as potential prognostic indicators in melanoma patients. Sci. Rep. 2021, 11, 1023. 10.1038/s41598-020-80336-8

21. Lim, Y.; Ku, N.O. Revealing the roles of keratin 8/18-associated signaling proteins involved in the development of hepatocellular carcinoma. Int. J. Mol. Sci. 2021, 22, 6401. 10.3390/ijms22126401

22. Yan, J.; Yang, A.; Tu, S. The relationship between keratin 18 and epithelial-derived tumors: As a diagnostic marker, prognostic marker, and its role in tumorigenesis. Front. Oncol. 2024, 14, 1445978. 10.3389/fonc.2024.1445978

23. Rhee, H.; Kim, H.Y.; Choi, J.H.; Woo, H.G.; Yoo, J.E.; Nahm, J.H.; Choi, J.S.; Park, Y.N. Keratin 19 expression in hepatocellular carcinoma is regulated by fibroblast-derived HGF via a MET-ERK1/2-AP1 and SP1 axis. Cancer Res. 2018, 78, 1619–1631. 10.1158/0008-5472.CAN-17-0988

24. Xie, J.; Guo, T.; Zhong, Z.; Wang, N.; Liang, Y.; Zeng, W.; Liu, S.; Chen, Q.; Tang, X.; Wu, H.; Zhang, S.; Ma, K.; Wang, B.; Ou, Y.; Gu, W.; Chen, H.; Qiu, Y.; Duan, Y. ITGB1 drives hepatocellular carcinoma progression by modulating cell cycle process through PXN/YWHAZ/AKT pathways. Front. Cell Dev. Biol. 2021, 9, 711149. 10.3389/fcell.2021.711149

25. Zhang, Z.; Peng, Z.; Cao, J.; Wang, J.; Hao, Y.; Song, K.; Wang, Y.; Hu, W.; Zhang, X. Long noncoding RNA PXN-AS1-L promotes non-small cell lung cancer progression via regulating PXN. Cancer Cell Int. 2019, 19, 20. 10.1186/s12935-019-0734-0

